# Astrocytes derived from ASD patients alter behavior and destabilize neuronal activity through aberrant Ca^2+^ signaling

**DOI:** 10.1101/2021.10.11.463231

**Authors:** Megan Allen, Ben S. Huang, Michael J. Notaras, Aiman Lodhi, Estibaliz Barrio Alonso, Paul Wolujewicz, Jonathan Witztum, Francesco Longo, Maoshan Chen, David Greening, Eric Klann, M. Elizabeth Ross, Conor Liston, Dilek Colak

**Affiliations:** Center for Neurogenetics, Feil Family Brain and Mind Research Institute, Weill Cornell Medicine, Cornell University, New York, USA; Center for Neural Science, New York University, New York, USA; Molecular Proteomics Laboratory, Baker Heart & Diabetes Institute, Melbourne, Victoria, Australia; Gale & Ira Drukier Institute for Children’s Health, Weill Cornell Medicine, Cornell University, New York, USA; Department of Psychiatry, Weill Cornell Medicine, Cornell University, New York, USA

**Author notes:** These authors contributed equally to this work (Megan Allen, Ben S. Huang).

**Keywords:** Astrocytes, Calcium signaling, chimeric mice, Induced pluripotent stem cells, ASD, Autism, and Cell transplantation

## Abstract

The cellular mechanisms of Autism Spectrum Disorder (ASD) are poorly understood. Cumulative evidence suggests that abnormal synapse function underlies many features of this disease. Astrocytes play in several key neuronal processes, including the formation of synapses and the modulation of synaptic plasticity. Astrocyte abnormalities have also been identified in the postmortem brain tissue of ASD patients. However, it remains unclear whether astrocyte pathology plays a mechanistic role in ASD, as opposed to a compensatory response. To address this, we strategically combined stem cell culturing with transplantation techniques to determine disease specific properties inherent to patient derived astrocytes. We demonstrate that ASD astrocytes induce repetitive behavior as well as impair memory and long-term potentiation when transplanted into the healthy mouse brain. These *in vivo* phenotypes were accompanied by reduced neuronal network activity and spine density caused by ASD astrocytes in hippocampal neurons *in vitro*. Transplanted ASD astrocytes also exhibit exaggerated Ca^2+^ fluctuations in chimeric brains. Genetic modulation of evoked Ca^2+^ responses in ASD astrocytes modulates behavior and neuronal activity deficits. Thus, we determine that ASD patient astrocytes are sufficient to induce repetitive behavior as well as cognitive deficit, suggesting a previously unrecognized primary role for astrocytes in ASD.

## Introduction

ASD is a developmental disability characterized by impaired social communication, restrictive repetitive behaviors, and quite often, cognitive deficits. Nearly 95% of ASD diagnoses are idiopathic or not associated with a known genetic mutation(s)^1^. Our lack of knowledge about the cell types that participate in disease progression early in brain development has limited the generation of effective therapies. While ASD is highly heritable (40-80%), uncovering common genetic biomarkers through sequencing and linkage studies remains challenging^2–4^. This suggests that a multitude of rare genetic variants converge on limited biological pathways to cause ASD. Recent work has begun to cluster these rare variants and indicate that most play a role in neural communication and plasticity^2, 5, 6^.

Intriguingly, the onset of ASD symptoms coincides with activity-dependent synaptic plasticity events^7^. In typically developing children, intense neuroplasticity occurs in the first few years of life, a time which parallels significant astrocyte proliferation and refinement^8^. In fact, astrocytes are in close physical contact with neuronal cell bodies, dendrites, and dendritic spines. A single mouse cortical astrocyte contacts over 100,000 synapses, whereas a human astrocyte contacts up to 2,000,000 synapses^9–11^. This intimate physical relationship supports the instructive roles astrocytes play in several key plasticity processes, including the formation of synapses^12–14^, secretion of factors that affect enhance structural changes like spine formation and dendritic arborization^15–18^, pruning of supernumerary synapses^19^, and the removal of excess neurotransmitters to prevent excitotoxicity^20^. Further, astrocytes have been implicated in the maintenance of learning and memory through the modulation of long-term potentiation (LTP)^21^. LTP, one of the major forms of synaptic plasticity, represents the cellular basis for learning and memory^22^. However, whether human astrocyte pathology plays a causal role in nonsydromic ASD, as opposed to a compensatory response to an already diseased brain, is not clear.

Astrocytes have been implicated in the pathogenesis of mouse models of syndromic ASDs where a single mutation leads to disease development. Co-culture experiments revealed that the presence of astrocytes derived from the brains of two syndromic mouse models, Rett and fragile X syndromes, negatively altered neuronal structure, which impeded proper synaptic transmission^23–24^. Additionally, transcriptomic and immunohistochemistry studies in postmortem tissue found enrichment of astrocytic reactivity in the cortex of brains from individuals with ASD^25–27^. The first data demonstrating astrocyte dysfunction in nonsyndromic autism using induced pluripotent stem cell (iPSC) technology has recently been published^28^. In this study, patient iPSC-derived astrocytes caused a decrease in neurite number and synaptic markers in iPSC-derived neurons, suggesting a potential contribution of astrocyte dysfunction in ASD pathophysiology^28^. Despite this wealth of knowledge of astrocyte function in the healthy brain and implication of astrocyte dysfunction in synaptic deficits in ASD, specific behavioral alterations as well as mechanisms through which astrocytes contribute to ASD remain unknown.

Here, we show that the presence of ASD astrocytes in a healthy brain induces repetitive behavior as well as memory and synaptic plasticity deficits. Three independent experiments, unbiased proteomic analysis, *in vitro* two-photon live-cell imaging, and live-animal two-photon imaging, commonly identify that altered Ca^2+^ signaling is an inherent defect in ASD astrocytes derived from multiple patients. We subsequently combined patient stem cell culturing methods with transplantation techniques to generate chimeras to study how astrocytes derived from ASD patients’ iPSCs function *in vivo*. Of note, in a physiological environment, transplanted patient astrocytes exhibit elevated Ca^2+^ responses. This phenotype is also recapitulated in cultured patient astrocytes. We report that ASD astrocytes induce repetitive behavior as well as memory impairments, which are accompanied by reduced LTP in hippocampal slice cultures from ASD chimeras. In line with these phenotypes, ASD astrocytes reduce spine density and neuronal network activity when co-cultured with wild type (WT) hippocampal neurons *in vitro*. Modulation of evoked Ca^2+^ release via genetic manipulation of IP_3_ receptors (IP_3_Rs) in ASD astrocytes protects against deficits in neuronal network dynamics and associative memory behavior. In summary, these data define a mechanistic role for astrocytes in ASD, revolutionizing our understanding of ASD pathogenesis.

### Aberrant Ca^2+^ activity in ASD astrocytes

To obtain ASD patient astrocytes, we used iPSCs derived from healthy control subjects (CTRL) and ASD patients. In total, we employed 9 individual CTRL (7 male, 2 female) and 9 individual ASD (all male) iPSC lines, most of which are deposited at NIMH and CIRM repositories. Only two CTRL lines come from the Coriell Institute. Please refer to **Supp Table 1** for details about patient clinical information, ADOS scores, age of sampling, race, etc. as well as identifier number for purchasing information. We performed Whole Exome Sequencing (WES) in astrocytes derived from all CTRL and ASD lines (18 in total) used in this study. In our WES analyses, in which we comprehensively interrogated single nucleotide variants (SNVs) and indels as well as copy number variants (CNVs), we did not find any recurrent SNV or CNV known to be associated with ASD in either our cases or controls. Additionally, individuals with ASD did not have a higher burden of rare exonic CNVs altering autism associated genes compared to controls, thereby supporting the clinical implication that the lines for our cases come from nonsyndromic ASD patients. **Supp Table 2** summarizes the counts of hits for LGD SNVs and LGD CNVs and **Supp Table 3** provides the information of variants in ASD cases and which individual carries them.

To obtain ASD patient astrocytes without biasing differentiation, we integrated two distinct strategies for successful generation of human iPSC-derived astrocytes^29, 30^. 3D cortical spheroids spontaneously generate astrocytes, which express similar protein profiles as purified human brain astrocytes^29, 31^. Here, we adapted an undirected-differentiation organoid system^32, 33^ to isolate astrocytes derived from CTRL or ASD patient lines (**Fig. 1a** and **Supp Fig. 1a**). Because this organoid protocol relies on spontaneous generation of cell types, it likely preserves disease-specific signatures unique to patient astrocytes. Undirected organoid protocols recapitulate the temporal sequence of cortical development seen in developing embryos^34^, and so, patient organoids spontaneously generate astrocytes in an environment that mimics the early ASD brain. Specifically, ASD astrocytes develop in the presence of genetically matched ASD neurons. This is important because astrocytes require early interaction with neurons to induce expression of critical receptors and trigger temporally regulated activation patterns^35^. To expand CTRL and ASD astrocytes in 2D culture, we enzymatically dissociated organoids at 75 days *in vitro* (DIV). We next selected for astrocyte enrichment with cell culture medium supplemented with glucose and low serum content (2%) as previously described^30^. While serum might select for specific astrocytic phenotypes, it is required for astrocyte expansion. Astrocytes grown in our selection medium expressed numerous astrocyte markers including ALDH1L1, GFAP, and AQP4 (**Fig. 1b**) as well as Vimentin and S100Beta (**Supp Fig. 1**b, c) by passage 8. Additionally, these astrocytes displayed fairly weak F-actin signal, whose levels are positively correlated with reactive phenotypes, indicating that CTRL and ASD astrocytes were not reactive when in culture (**Supp Fig. 2**).

**Fig. 1:**
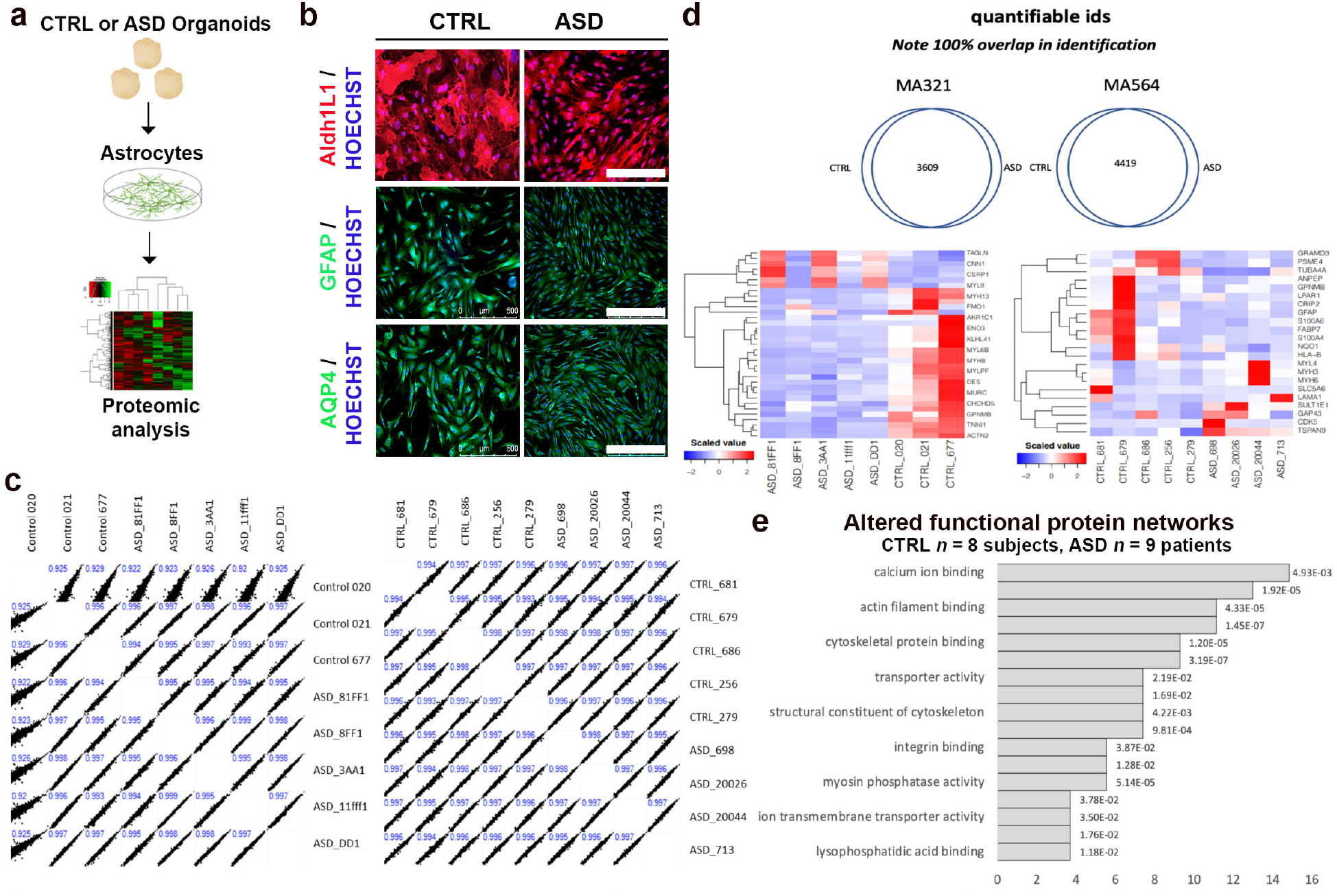
Aberrant Ca^2+^ signaling in ASD astrocytes. **a,b**, Spontaneous generation of human astrocytes. **a,** We dissociated astrocytes from ASD or CTRL organoids at day 75 and expanded them in culture (see also Methods section). **b**, Representative images from immunostainings shows that astrocytes dissociated from organoids expressed multiple astrocyte markers: ALDH1L1, GFAP, and AQP4 as well as Vimentin and S100Beta (see also **Supp** Fig. 1b-d). **c-e**, Proteomic study identified Ca^2+^ signaling as the most significantly altered network in ASD astrocytes. Proteins were extracted from astrocytes and labeled with Tandem Mass Tag (TMT) chemistry followed by LC/MS analysis. **c**, Two independent proteomic runs and analyses, of which each exhibited high experimental reproducibility. **d**, Venn diagrams for both runs revealed 3609 and 4419 proteins, respectively, which were common to both CTRL and ASD astrocyte samples. **e**, GO analysis revealed enrichment for Ca^2+^ ion binding proteins in ASD samples. **This experiment suggests that ASD astrocytes harbor dysfunctional Ca^2+^ signaling.** Scale bar = 500 μm. Data are represented as mean ± SEM. Proteomics analysis: CTRL *n* = 8 lines; ASD *n* = 9 lines.

To identify the inherent pathological properties within ASD astrocytes, we first took an unbiased approach and compared the protein profiles of 9 ASD astrocytes to 8 healthy CTRL astrocytes with Tandem mass tag (TMT) liquid chromatography mass spectrometry (**Fig. 1c-e**)^36^. Notably, proteomic analysis validated our method of astrocyte extraction as several markers for astrocyte identity were detected (**Supp Fig. 3**)^37, 38^. Expression of Vimentin, ERBB2, Na^2+^/K^+^ ATPase, and LGALS3 in the proteomic profile of our cultured astrocytes (**Supp Fig. 3**a-d) suggested that sufficient neuronal astrocyte interaction occurred in the organoid system.

Proteomics analysis was split into two label-based protein quantification analyses using Tandem-Mass-Tag (TMT) chemistry. This approach elicited a median depth of 4014 protein quantifications (Run 1: 3609, Run 2: 4419). Quantifiable values were also highly reproducible between samples and runs (CTRL *r*^2^ = 0.94-0.995; ASD *r*^2^ = 0.995-0.996; **Fig. 1c**). This demonstrates high experimental reproducibility and cell-type homogeneity. For each sample group, differentially regulated proteins constituted 22 (0.969% of Run 1) and 35 proteins (0.497% of Run 2) of each analysis group, respectively (**Fig. 1d**). Higher magnification of heat maps and individual gene names are presented in **Supp Fig. 3**e and f, respectively. We also screened 12 specific *reactive* astrocyte markers^39, 40^, 9 of which were not detected and 3 were low abundance within our astrocyte proteomics datasets (see **Supp Fig.3** legends). Together with the observation of fairly low F-actin expression in these cells (**Supp Fig. 2**), this data suggest that our astrocytes are not in an overtly reactive state.

Pathway enrichment analysis for categorical data was performed based on a Fisher’s exact test with a Benjamini–Hochberg FDR threshold of 0.02. Coverage and enrichment in Molecular Function (MF) GO categories revealed Ca^2+^ ion binding as the most enriched MF category (**Fig. 1e**). Further to this, Qiagen’s Ingenuity Pathway Analysis (IPA) was adapted to compare the two independent runs, and once more determined Ca^2+^ signaling as the most significantly altered pathway. These results collectively suggest that ASD astrocytes might exhibit aberrant Ca^2+^ signaling.

Taken together, these data suggest that ASD astrocytes respond to stimulation with increased Ca^2+^ responses.

### Organoid-derived astrocytes migrate throughout the mouse cortex and survive into adulthood

To determine whether ASD astrocytes induce behavioral deficits, we transplanted CTRL or ASD astrocytes into the brains of neonatal mice using an established multisite-injection protocol with minor modifications^42^. We obtained astrocytes from organoids and expanded in 2D culture as described above. Prior to transplantation, we transduced astrocytes with a CAG-GFP lentivirus to distinguish human derived cells from endogenous mouse cells *in vivo*^43^. CTRL and ASD astrocytes were enzymatically dissociated into a single cell solution and subsequently transplanted into the brains of neonatal *Rag2^KO^* immune-compromised mice at postnatal days 1-3 (P1-3). We used *Rag2^KO^* mice as transplant hosts to limit rejection of human astrocytes derived from multiple individuals^42, 44, 45^. We transplanted a total of 8-10x10^5^ astrocytes into each mouse brain, spread out over four injection sites, which were bilateral along the midline, anterior and posterior to bregma.

This transplantation technique allowed for broad and homogenous human cell distribution throughout the cortex and hippocampus (**Fig. 2**). Notably, we did not find gross differences in the migratory patterns of GFP+ human cells in mouse brains that were collected at P60. Representative images from whole brain sagittal sections highlighted extensive migration throughout the cortex and subcortical regions (**Fig. 2b**). In addition to GFP labeling, we also confirmed astrocyte survival and migration in chimeric brains at P60 by immunostaining against a human specific GFAP epitope^46^ (**Fig. 2c-e**). To quantify the number of human astrocytes that survived and incorporated in the chimeric brains, we performed stereological cell counting using the optical fractionator method. This method uses systematic random sampling to obtain a statistically valid sample of the region of interest, thereby allowing unbiased analysis. We did not find a significant difference in the number of surviving human astrocytes in ASD astrocyte chimeric brains relative to CTRL astrocyte chimeric brains (**Supp Fig. 4**). To determine whether astrocytes maintained their identities upon transplantation and maturation, we co-immunostained chimeric brains for GFP and either GFAP or ALDH1L1 (red), two well described astrocyte markers (**Fig. 2f,g**). More than 90% of human GFP+ cells co-expressed astrocyte markers in adult chimeric brains (**Fig. 2h,i**). Of note, we partially automated cell counting using ImageJ, and in doing so, applied a stringent thresholding protocol (which may have prevented detection of human GFP+ astrocytes that expressed either ALDH1L1 or GFAP at low levels). We also co-immunostained chimeric brains for GFP and neuron marker NeuN. We examined over 800 GFP+ cells in 9 chimeric brains (transplanted with 4 distinct iPSC lines/4 CTRL subjects and 5 distinct iPSC lines/5 ASD patients) and found that none of the GFP+ cells expressed nuclear NeuN or exhibited a typical neuronal morphology (**Supp Fig. 5**). This is consistent with the fact that organoid-derived cultured cells had already acquired an astrocyte fate prior to injections and were transplanted following completion of neurogenesis when astrocyte generation innately peaks within the postnatal brain.

**Fig. 2:**
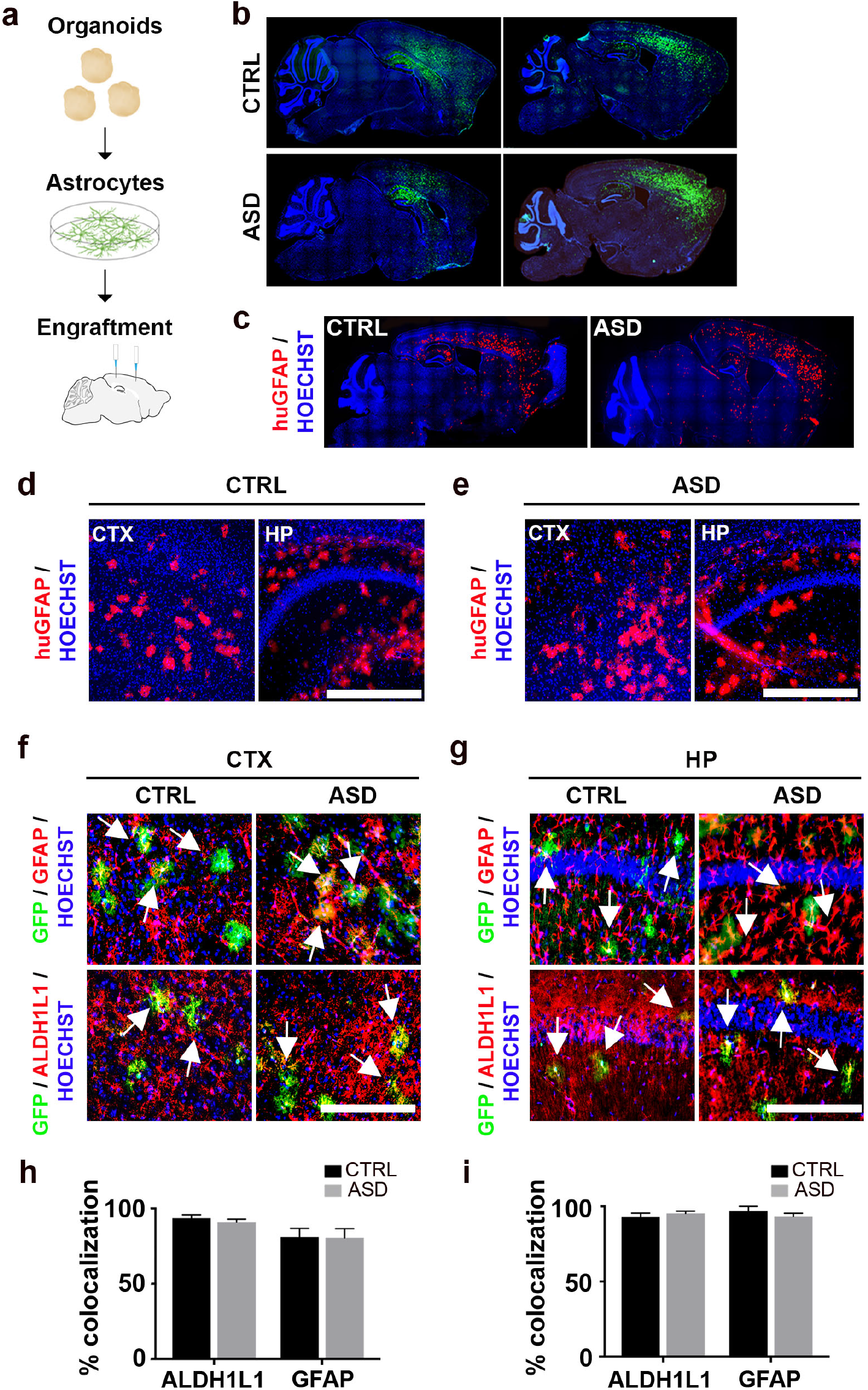
Organoid-derived astrocytes migrate throughout the mouse cortex and survive into adulthood. **a**, Schematic of experimental workflow. We transplanted a total of 8-10x10^5^ astrocytes (infected with CAG-GFP virus) into the brains of *Rag2^KO^* neonatal mice, spread out over four injection sites, which were bilateral along the midline, anterior and posterior to bregma. **b**-**e**, immunostained whole brain slices cut on the sagittal plane both for GFP (**b**) and the human-specific GFAP epitope (huGFAP) at P60 showed a wide and homogenous spread throughout the cortex (**c**) (see also **Supp** Fig. 4 for stereological quantifications). **d,e**, Representative higher magnification images illustrated the spread of huGFAP in cortex and hippocampus of chimeric mice. **f**-**i**, Representative co-immunostained images of chimeric brains revealed high co-localization of GFP expression (human astrocytes, green) with the two astrocyte markers, GFAP (red, top panel) and ALDH1L1 (red, bottom) in the cortex (**f**) and hippocampus (**g**). Arrows indicate examples of dual positive cells. More than 90% of GFP+ cells expressed astrocyte markers suggesting that astrocytes retained their identities upon maturation in the adult mouse brain (**h** and **i**, see also **Supp** Fig. 5). **These results establish that CTRL and ASD astrocytes dissociated from organoids generated homogenous transplantations in host brains.** CTX: Cortex; HP: Hippocampus. Scale bar = 500 μm for d and e and 250 μm for f and g. Data are represented as mean ± SEM. Co-localization: CTRL and ASD *n* = 8 per group (2 mice/2 distinct lines, and 4 slices per brain).

### *In vivo* imaging confirms aberrant Ca^2+^ activity in ASD astrocytes

Because Ca^2+^ transients both in soma and branches contribute to Ca^2+^ signaling in astrocytes, astrocyte Ca^2+^ transients are best captured *in vivo* where astrocytes display their mature and complex morphology^47^. To validate our *in vitro* findings in an *in vivo* system, we used two-photon microscopy to image Ca^2+^ activity in transplanted human astrocytes in awake behaving mice responding to an environmental stimulus. To measure Ca^2+^ responses in human astrocyte chimeric mice, we used genetically encoded indicators and cranial window implantation followed by two-photon imaging as previously described^48^ (**Fig. 3a-c**). We virally transfected the astrocytes with AAV carrying the genetically encoded calcium indicator GCaMP6f (AAV2/5-*GfaABC_1_D-GCaMP6f*). We engrafted the GCaMP6f-expressing human astrocytes into newborn *Rag2*^KO^ (P1-3) by injecting the cells into the frontal cortex. At P60+, we implanted a 3-mm-diameter glass cranial window over the primary somatosensory and motor cortices (S1/M1), attached a titanium head-bar to the skull for head-fixation, and secured the implants with dental cement (see **Methods** for further details). We allowed the mice to recover post-surgery for at least 3 weeks. Finally, we performed *in vivo* two-photon imaging to record Ca^2+^ activity in transplanted human astrocytes within live mice actively responding to the environment. We segmented the cells with a 20-µm-diameter mask, which encompassed both soma and processes. Transplanted human astrocytes displayed a variety of Ca^2+^ activity responses *in vivo* in reaction to an air-puff startle stimulus, including increased, decreased, and unchanged Ca^2+^ levels (**Fig. 3d**). When categorized, these response groups revealed significant differences between Ca^2+^ activity in CTRL and ASD astrocytes. In the increased response group, ASD astrocytes exhibited a significantly larger increase than CTRL cells (**Fig. 3e**,top). While decreases in Ca^2+^ levels were not significant between the groups, ASD astrocytes displayed a drastic increase in Ca^2+^ fluctuations compared to the CTRL astrocytes in decreased Ca*^2+^* response subgroup (**Fig. 3e**,middle). Similarly, while CTRL cells showed a completely flat and unresponsive activity profile, the ASD cells with unchanged Ca^2+^ levels displayed a highly significant change in Ca^2+^ fluctuations (**Fig. 3e**,bottom).

**Fig. 3:**
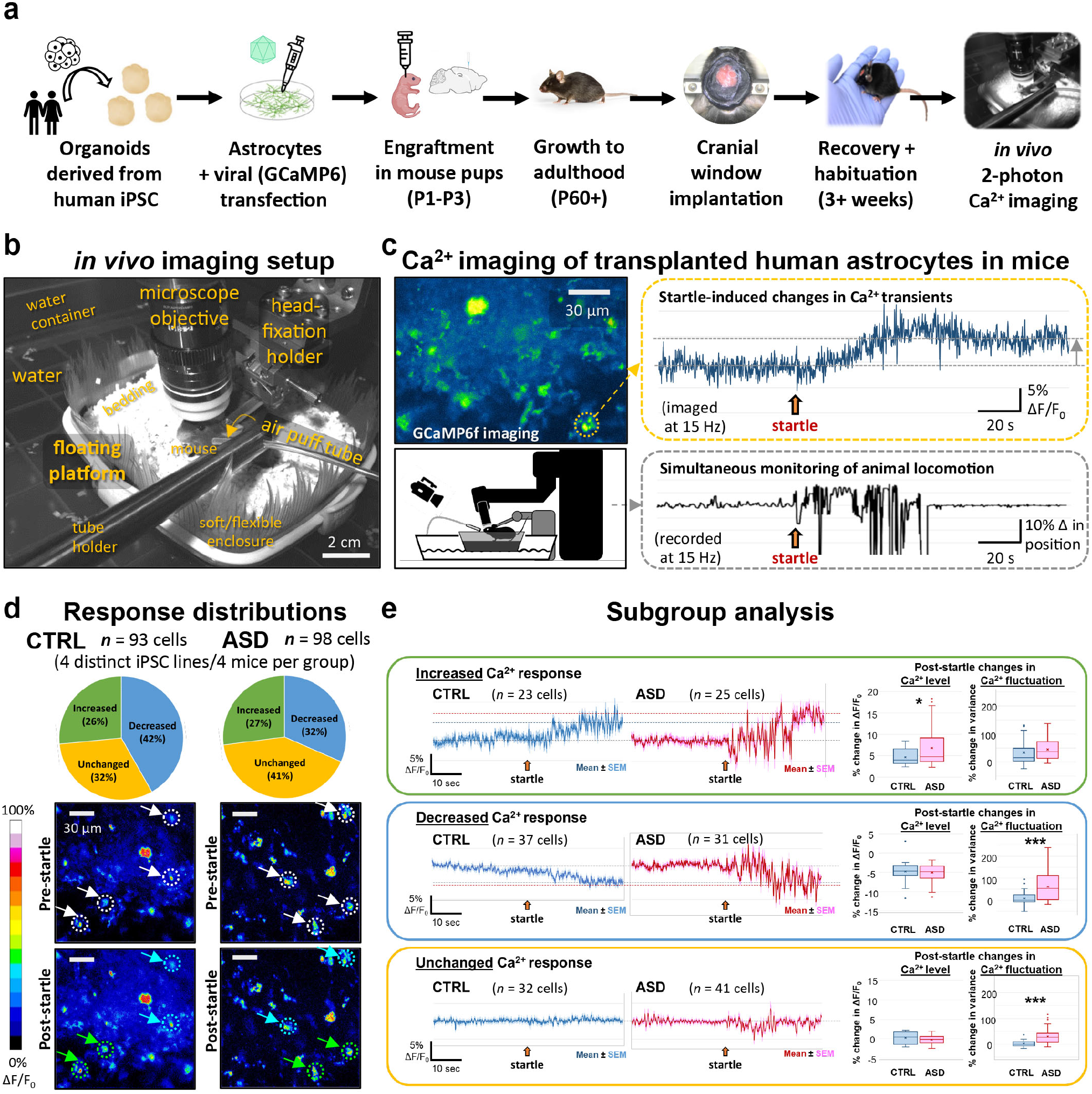
*In vivo* imaging confirms aberrant Ca^2+^ activity in ASD astrocytes. **a**, Schematic of experimental workflow. **b,** *In vivo* imaging setup. Photograph with labeled components of our custom-designed floating platform, developed to provide a tactile virtual-reality environment for head-fixed mice and to enable imaging of actively locomoting animals with minimal motion confounds (see **Methods** for details). **c,** Ca^2+^ imaging of transplanted human astrocytes in mice. *Left, upper panel*: a representative image of GCaMP6f-expressing human astrocytes in the cortex of a mouse engrafted with cells derived from a CTRL human subject. *Right, upper panel (orange box)*: Ca^2+^ transient recorded from the cell marked by the orange circle on the left image, showing an increase in Ca^2+^ level after the startle/air-puff stimulus (*orange arrow*). *Left bottom panel*: Schematic of the imaging setup. The mouse is situated at the center on top of an enclosed platform (boat) floating on water (see **Methods** for details). *Right bottom panel (gray box)*: Representative trace of recorded locomotion during imaging, produced by analyzing the infrared video recording and plotting the light intensity change of a select ROI (region of interest) on the floating platform. As animal movement directly translated into platform displacement, any body motion generated by the animal could be tracked with accuracy and high temporal resolution, even heavy breathing could be seen indicated by the brief ticks at the latter part of the recording. **d,** Response distributions. Transplanted human astrocytes displayed a variety of Ca^2+^ activity responses *in vivo* in reaction to the air-puff startle stimulus, including increased, decreased, and unchanged Ca^2+^ levels. We segmented the cells with a 20-µm-diameter mask, which encompassed both soma and processes. *Top*: Pie charts showing the percent distribution of each response types in CTRL and ASD astrocytes. *Bottom*: Representative images of CTRL and ASD astrocytes before and after startle. *White circles and arrows* (*upper panels*) point to sample cells showing post-startle increased (*green arrows/circles*) and decreased (*blue arrows/circles*) Ca^2+^ responses. **e,** Subgroup analysis. For all traces, the mean ± SEM are plotted, representing the population average. *Top (green box)*: Increased response (cells displaying a positive change in ΔF/F after startle). Box plots showing ASD cells exhibit a significantly larger increase than CTRL cells (post-startle % change in ΔF/F: CTRL: +4.66 ± 0.384%, *n* = 23; ASD: +6.81 ± 0.937%, *n* = 25; unpaired *t*-test, *p* = 0.04). No significant differences are found in the changes in fluctuation between CTRL and ASD cells. *Middle (blue box)*: Decreased Ca^2+^ response subgroup (cells showing negative change in ΔF/F after startle). Both CTRL and ASD show decreases in Ca^2+^ to similar levels (∼ -5%). However, there is a drastic difference in their change in Ca^2+^ fluctuations. Compare the flat downward sloping blue trace vs. the red trace with the dramatic fluctuation after startle. The box plots quantitatively illustrate this difference (Post-startle change in Ca^2+^ fluctuation: CTRL: 7.08 ± 4.74%, *n* = 37; ASD: 60.74 ± 11.04%, *n* = 31; unpaired *t*-test, *p* =0.000062). *Bottom (orange box)*: Unchanged Ca^2+^ response subgroup (cells whose changes in Ca^2+^ levels are between +2 and -2%). While CTRL cells in this group show a completely flat and unresponsive activity profile, the ASD cells with unchanged Ca^2+^ levels showed a highly significant change in Ca^2+^ fluctuation (30% increase in variance) (Post-startle change in Ca^2+^ fluctuation: CTRL: 1.39 ± 2.08%, *n* = 32; ASD: 30.75 ± 4.63%, *n* = 41, unpaired *t*-test, *p* = 0.00000036).

Together, *in vivo* Ca^2+^ imaging of transplanted human astrocytes in awake behaving mice indicate that ASD astrocytes exhibit significantly heightened responses to environmental stimuli, as seen both in significantly increased Ca^2+^ elevation as compared to CTRL astrocytes and highly significant increases in all ASD astrocytes in response to the startle stimulus despite decrease or unchanged Ca^2+^ levels.

### ASD astrocyte chimeric mice exhibit repetitive behavior as well as impaired memory and hippocampal LTP

We next sought to assess whether ASD astrocyte chimeric mice display cognitive and behavioral abnormalities. We subjected adult CTRL and ASD chimeric mice (between 3 and 5 months of age) to a range of ethologically relevant behavioral assays. We first evaluated general locomotor activity and exploratory behavior in chimeric mice with an automated Open Field test. ASD chimeric mice did not demonstrate differences in exploratory or general activity relative to CTRL chimeric mice (unpaired *t* test, *p* value = 0.782). These results also suggest that our injection protocol did not affect ambulatory activity and is not a confounding factor in the interpretation of other behavioral results (**Supp Fig. 6**).

To assess learning and memory in ASD and CTRL astrocyte chimeric mice, we used a classical fear conditioning protocol (**Fig. 4a**). Fear conditioning is an associative learning and memory task where mice are trained to associate a neutral conditioned stimulus (CS; audible tone, 70dB) with an aversive unconditioned stimulus (US; mild electrical foot shock, 0.7mA) and display a conditioned response (CR; freezing behavior). The freezing behavior is used as an index of the mouse’s ability to learn the task and later recall the associative memory. On day one, mice were exposed to a 5-min conditioning trial that involved three co-terminating tone shock pairings. We evaluated acquisition of the association by plotting the percentage of time spent freezing (% freezing) with each successive pairing. On day two, mice were returned to the conditioning context for 5 min, with % freezing measured for the first 2 min of the test session. This measured the association of the memory to the context, which is hippocampal dependent. On day three, we placed mice in a novel context and presented only the audible tone. Freezing behavior to the first tone presentation was recorded and represents cue-associated memory. While there was no significant difference between CTRL and ASD astrocyte chimeric mice in their ability to learn the association between tone and shock during the acquisition phase (**Fig. 4b**), ASD astrocyte chimeric mice exhibited impaired contextual memory relative CTRL astrocyte chimeric mice (**Fig. 4c,d**). We also assessed spatial learning and memory in ASD astrocyte chimeric mice. To do this, we used the Morris water maze, where a test mouse relies on distal cues to locate a hidden escape platform submerged in opaque water. Spatial learning is acquired though repeated trials and memory is determined through efficient navigation to an area that formerly housed the escape platform. No spatial learning and memory deficits were detected in ASD astrocyte chimeric mice in the Morris water maze (**Supp Fig. 7**a-d).

**Fig. 4:**
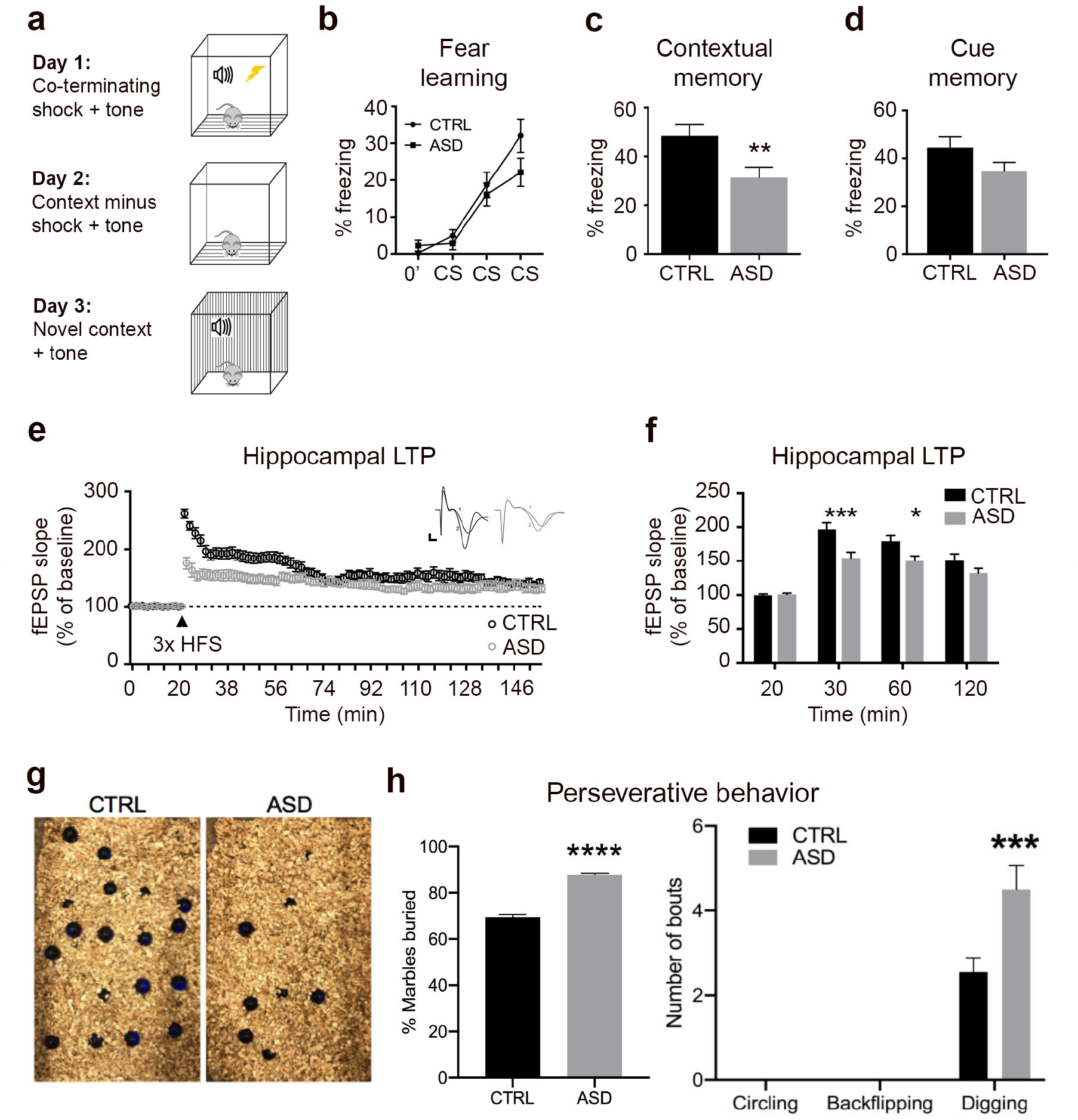
ASD astrocyte chimeric mice exhibit impaired associative memory and hippocampal LTP along with repetitive behavior. **a-d**, ASD astrocyte chimeric mice displayed deficits in associative memory but not in associative learning. **a**, Schematic summarizing classical fear conditioning paradigm. On day 1, mice were trained to associate an audible tone (30 sec duration, 70dB) with a co-terminating foot shock (1-sec duration, 0.7mA). Testing days 2 and 3 measured freezing behavior in response to exposure to the training context or an audible cue in a novel context, respectively. Freezing behavior in the testing trials provided a quantifiable measure of associative memory (see also Methods). **b**, There was no significant difference in the rate of acquisition learning between CTRL and ASD mice (ANOVA with Bonferroni posttests *p* value > 0.05). **c**, ASD chimeric mice showed reduced freezing behavior when exposed to the fear context (unpaired *t* test = 0.009). **d**, No differences were found in the freezing behavior between CTRL or ASD chimeric mice during cue presentation in a novel context (unpaired *t* test = 0.10). **e,f,** We tested LTP as a synaptic correlate of learning and memory in hippocampal brain slices (400 µm) of 4-6-month old transplanted mice. ASD astrocyte chimeric brain slices showed reduced potentiation in the initial phase of LTP compared to control (30-60 mins). ASD astrocyte chimeric brain slices displayed the greatest differences in the fEPSP slope within the first 60 minutes of recording (**f**). **g**,**h,** To test ASD chimeric mice for repetitive behavior, we employed the marble burying test as well as monitored circling and backflipping. Marbles were scored as buried if at least 60% of the marble was covered (30-min period) (**g**). ASD astrocyte chimeric mice buried significantly more marbles when compared to CTRL astrocyte chimeric mice (unpaired *t* test = 0.009), but exhibited no circling or backflipping (**h**). **Taken together, these results indicate that ASD astrocytes induce ASD-related perseverative behavior, memory dysfunction, and hippocampal LTP deficits.** Data are represented as mean ± SEM. Fear testing: CTRL *n* = 21 male and female mice/4 distinct lines, ASD *n* = 22 male and female mice/3 distinct lines; LTP: CTRL *n* = 10 slices, 4 mice/2 distinct lines; ASD *n* = 10 slices, 5 mice/2 distinct lines). Repetitive behavior: CTRL *n* = 16 mice/5 distinct lines, ASD *n* = 13 mice/5 distinct lines.

Persistent changes in synaptic strength via LTP represent a cellular mechanism for the formation and retention of memories^37, 49^. Intriguingly, altered LTP has been implicated in multiple syndromic models of ASD and correlates with ASD relevant behavioral impairments^50^. Astrocyte support is not only required for proper synaptic plasticity^51–54^, but also enhances LTP and memory^55^. We, therefore, measured LTP in brain slices from CTRL and ASD astrocyte chimeric mice. We induced LTP with three 1-sec trains of 100 Hz stimulation with an intertrain interval of 60 sec in chimeric slices. We collected field excitatory postsynaptic potentials (fEPSPs) for an additional 140 min. ASD astrocyte chimeric slices showed reduced potentiation in the initial phase of LTP compared to CTRL astrocyte chimeric slices (**Fig. 4e,f**).

To assess ASD hallmark behavior, we evaluated both repetitive behavior and sociability. Per the DSM-5, ASD diagnosis is based on three categories of behavioral criteria: abnormal social interactions, communication deficits and repetitive behaviors^56^. We probed the role ASD astrocytes play in perseverative behaviors with a marble burying test^57, 58^. We administered marble burying in a cage that contained extra bedding at a depth of 5 cm with 28 marbles arranged in a 4x7 grid. Marbles that were at least 60% covered after a 30-min exploration period were scored as buried. ASD astrocyte chimeric mice buried significantly more marbles when compared to CTRL chimeric mice (unpaired *t* test = 0.009) indicating that ASD astrocyte transplantation induced a form of repetitive-like behavior (**Fig. 4g,h**). To score sociability, we used a three-chamber social interaction paradigm1^5, 59, 60^. In the sociability test, ASD and CTRL astrocyte chimeric mice showed no bias for any of the chambers during the habituation phase (**Supp Fig. 7**f). ASD astrocyte chimeric mice also spent similar amounts of time in the social and nonsocial zones relative to CTRL astrocyte chimeric mice (**Supp Fig. 7**g).

Results from these experiments indicate that ASD astrocytes can induce repetitive behavior as well as lead to selective memory impairments and altered synaptic plasticity.

### ASD astrocytes decrease neuronal network firing and spine density *in vitro*

To provide insight into mechanisms through which ASD astrocytes induce behavioral and LTP deficits in chimeric brains, we simulated the *in vivo* macroenvironment by co-culturing ASD astrocytes with mouse hippocampal neurons and assessed the effects of ASD astrocytes in structural plasticity and activity at a cellular level in these neurons. Hippocampal neuronal cultures are inherently a mixture of glutamatergic neurons (over 70%) and astrocytes (less than 30%) (**Supp Fig. 8**). We co-cultured primary hippocampal cells dissociated from wild type (WT) embryonic mouse brains between embryonic days 16 and 18 (E16-18) with human astrocytes isolated from CTRL or ASD organoids. These two co-culture conditions are the closest to our *in vivo* chimeric models in terms of cellular interactions. Because astrocytes mediate connectivity and synchronized network activity^61–63^, we first investigated whether ASD astrocytes disrupt neuronal network activity. At DIV14, we measured spontaneous network activity using multi-electrode array (MEA). In all co-culture experiments, we included an additional control culture dissociated from the same litter that consisted of primary hippocampal neurons and astrocytes but no human astrocytes (None). To assess network activity, we plated co-cultures on 48-well MEA plates with a 4:1 hippocampal cells to human astrocyte ratio (**Fig. 5a**). Each well contained 16 electrodes arranged in 4x4 grid that detected extracellular field potentials.

**Fig. 5:**
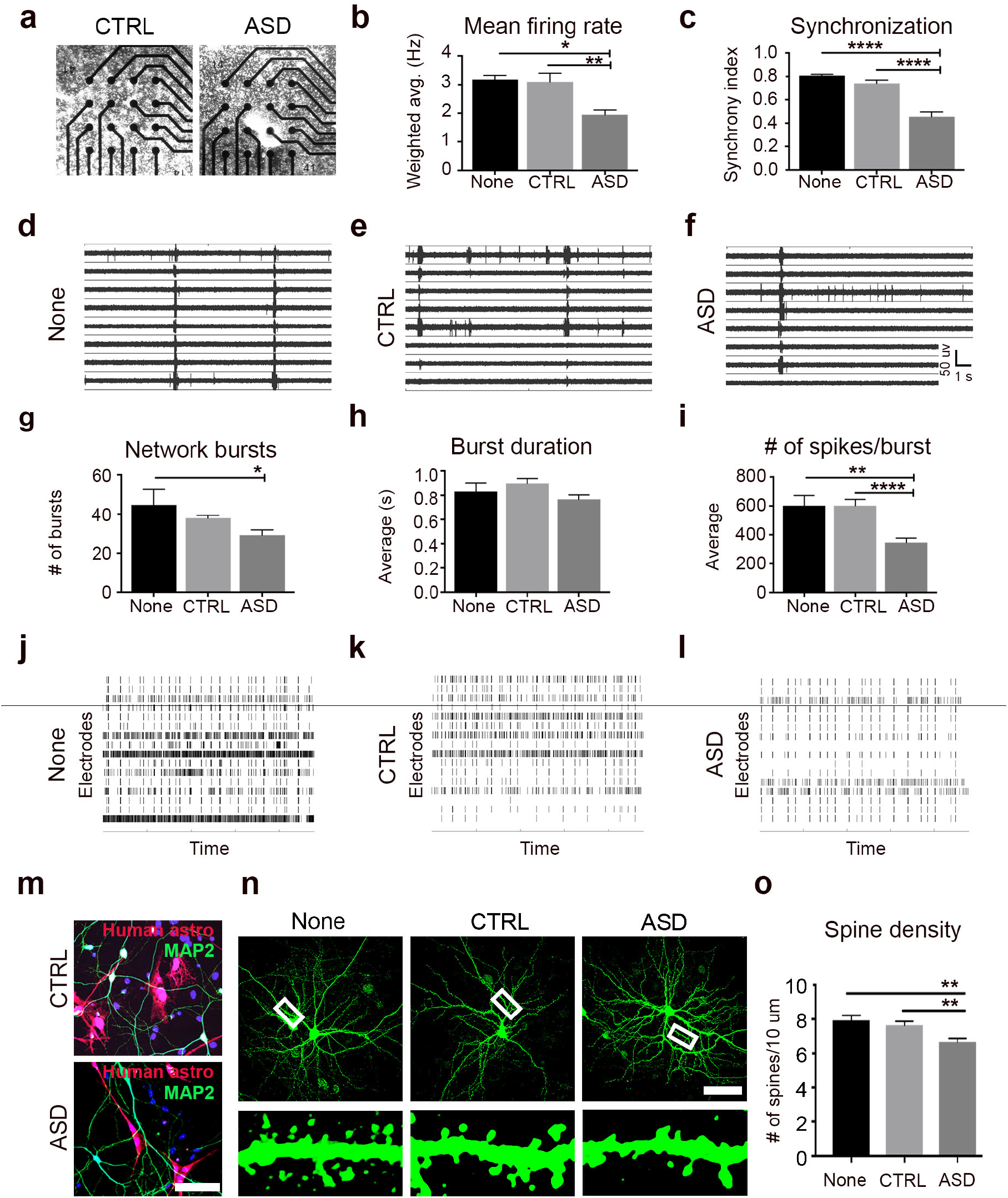
ASD astrocytes decrease neuronal network firing and spine density *in vitro*. **a-l**, We co-cultured primary hippocampal neurons dissociated from E16-18 WT mouse brains with human astrocytes isolated from CTRL or ASD organoids and measured spontaneous network activity with MEA. CTRL cultures refer to co-culturing of CTRL human astrocytes with hippocampal neuronal cultures (naturally containing some mouse astrocytes), and ASD cultures refer to co-culturing of ASD human astrocytes with hippocampal neuronal cultures (naturally containing some mouse astrocytes). ‘None’ cultures refer to hippocampal neuronal cultures (containing some mouse astrocytes) without any addition of human astrocytes. **a**, Representative images of co-cultures plated in a single well of a 48-well array plate. **b**, ASD astrocyte co-cultures displayed decreased mean network firing rate when compared to CTRL co-cultures and None. (ANOVA with Tukey’s posttests None vs CTRL *p* value = 0.98, None vs. ASD *p* value = 0.02, CTRL vs. ASD *p* value = 0.002, see also **Supp Video 2**). **c**, ASD astrocytes also disrupted network synchronization when compared to CTRL astrocytes and None (None vs. CTRL *p* value = 0.65, None vs. ASD *p* value < 0.0001, CTRL vs. ASD *p* value < 0.0001). **d**-**f**, Representative raw traces of spontaneous spiking behavior over a 10-second period. **g**, ASD astrocytes decreased the # of network bursts when compared to None (None vs. ASD *p* value = 0.02, CTRL vs. ASD *p* value = 0.06). **h,i**, ASD astrocytes did not affect average burst duration but decreased the # of spikes per burst network (None vs. ASD *p* value = 0.005, CTRL vs. ASD *p* value < 0.0001). **j**-**l**, Representative raster plots (4-min) demonstrated the decrease in burst number and spikes per bursts in the ASD co-culture group compared to CTRL co-culture and None. **m**-**o**, Spine density quantified in a 10 μm dendritic segment at least 20 μm away from the soma in hippocampal neurons co-cultured with ASD or CTRL human astrocytes. **m**, Immunostained co-cultures with human astrocytes infected with CMV GFP lentivirus (pseudo colored red) prior to co-culture. **n**, Neurons were labeled with a αCamKII GFP AAV at DIV5 and fixed at DIV18 (10 μm dendritic segments shown at bottom). **o**, ASD astrocytes decreased spine density on primary hippocampal neurons (None vs. CTRL *p* value = 0.78, None vs. ASD 0.009, CTRL vs. ASD *p* value = 0.004). **These results provide direct evidence that ASD astrocytes influence structural and functional properties of neurons that weaken electrophysiological activity.** Scale bar = 100 μm. Data are represented as mean ± SEM. MEA: None *n* = 6 wells, CTRL *n* = 18 wells, 3 distinct lines; ASD *n* = 22-24 wells, 4 distinct lines. Spine quantification: None *n* = 25 neurons, CTRL *n* = 71 neurons co-cultured with 4 distinct lines, ASD *n* = 81 neurons co-cultured with 5 distinct lines. CTRL: Co-cultures of CTRL human astrocytes with mouse hippocampal neurons ASD: Co-cultures of ASD human astrocytes with mouse hippocampal neurons None: Mouse hippocampal neuron cultures with no human astrocytes

We evaluated the following properties of spontaneous network firing in our co-cultures: mean firing rate, synchronicity, and network bursting behavior. Addition of ASD astrocytes to hippocampal neurons decreased the mean firing rate compared to the neurons co-cultured with CTRL human astrocyte and None (mouse hippocampal cultures with no human astrocytes) (**Fig. 5b**, see also **Supp Video 2**). ASD astrocytes also disrupted synchronous firing (**Fig. 5c**). Representative raw spike plots further illustrated the diminished mean firing and reduced synchronicity in the ASD co-cultures relative to CTRL co-cultures and None (**Fig. 5d-f**). We also assessed bursting behavior, which is important for information transfer across brain. Network bursts occur when groups of neurons fire coordinated trains of spikes in specified patterns. ASD astrocytes showed a decreased number of network bursts relative to None (**Fig. 5g**) and decreased number of spikes per burst compared to CTRL co-cultures and None (**Fig. 5i**). However, ASD astrocytes did not alter burst duration in these experiments (**Fig. 5h**). ASD co-cultures displayed reduced network bursts than our ‘None’ condition, but not our ‘CTRL’ cultures in **Fig. 5g**. Of note, while *p* value is 0.02 for None vs. ASD, it is 0.06 for CTRL vs. ASD. This strong trend suggests that this particular phenotype may only be subtle. Raster plots demonstrated the decrease in burst number and spikes per bursts in the ASD co-cultures (**Fig. 5j-l**). Overall, these experiments suggest that the addition of healthy human astrocytes into mouse primary neuronal cultures (already containing some mouse astrocytes) do not further support for network activity. However, we report that the addition of ASD astrocytes influences functional properties of neurons including mean firing rate, synchronization, and spike numbers.

To determine whether ASD astrocytes alter structural plasticity, we quantified spine density in neurons co-cultured with CTRL or ASD patient astrocytes or no human astrocytes (None) (**Fig. 5m**). To visualize dendrites and dendritic spines, we infected primary hippocampal neurons dissociated from E16-18 WT mouse brains with an αCaMKII-GFP AAV virus (**Fig. 5n**). We acquired maximum intensity projections and quantified spines along a 10 µm dendritic segment at least 20 µm away from the soma. The presence of CTRL astrocytes did not significantly affect the number of spines on neurons when compared to None. However, the presence of ASD astrocytes decreased spine density on neurons when compared to CTRL and None groups (**Fig. 5o**).

These experiments complemented the behavioral and plasticity phenotypes of chimeric mice in supporting the conclusion that ASD astrocytes induce specific cognitive and behavioral deficits by influencing the structural and functional properties of neurons.

### Modulation of Ca^2+^ responses in ASD astrocytes

We hypothesized that attenuating evoked intracellular Ca^2+^ responses in ASD astrocytes would prevent deficits in neuronal network function and behavior caused by ASD astrocytes with exaggerated evoked Ca^2+^ responses. Indeed, astrocytes respond to neuronal activity via elevated cytosolic Ca^2+^ concentrations^18, 64^. This triggers the intracellular signaling pathways that modulate neuronal connectivity. Thus, exaggerated evoked Ca^2+^ responses in ASD astrocytes could alter neuronal network activity and behavior.

To reduce evoked Ca^2+^ increases, we targeted inositol 1,4,5-trisphosphate receptors type 1 and 2 (IP_3_R) using shRNA based knockdown strategy (**Fig. 6**). IP_3_Rs are ligand gated calcium channels found on the surface of the endoplasmic reticulum (ER) and upon ligand binding, IP_3_Rs release Ca^2+^ that is stored in high concentrations within the ER. Canonical activation occurs in response to Gq linked G protein coupled receptor (GPCR) signal transduction^65^. Gq activators bind to a GPCR leading to the eventual liberation IP_3_R into the cytosol, which then binds to IP_3_Rs on the ER. Thus, reducing IP_3_R levels reduces Ca^2+^ release from the ER^66^ (see schematic in **Fig. 6a,b**). We exploited the known role of IP_3_R-mediated Ca^2+^ release to commonly lower evoked elevations in cytosolic Ca^2+^ in our patients’ astrocytes. Notably, because each patient line may exhibit varied mechanisms for increased cytosolic Ca^2+^, lowering IP_3_R levels would result in a common reduction in Ca^2+^ release from the ER across all patient-derived astrocytes. This approach therefore allowed for a universal modulation of Ca^2+^ signaling. We infected ASD astrocytes with a shRNA lentivirus that targeted IP_3_Rs (KD). As a negative control, we infected ASD or CTRL astrocytes with a non-targeting shRNA lentivirus (Non). The non-targeting control shRNA contained a sequence that did not target a gene product.

**Fig. 6:**
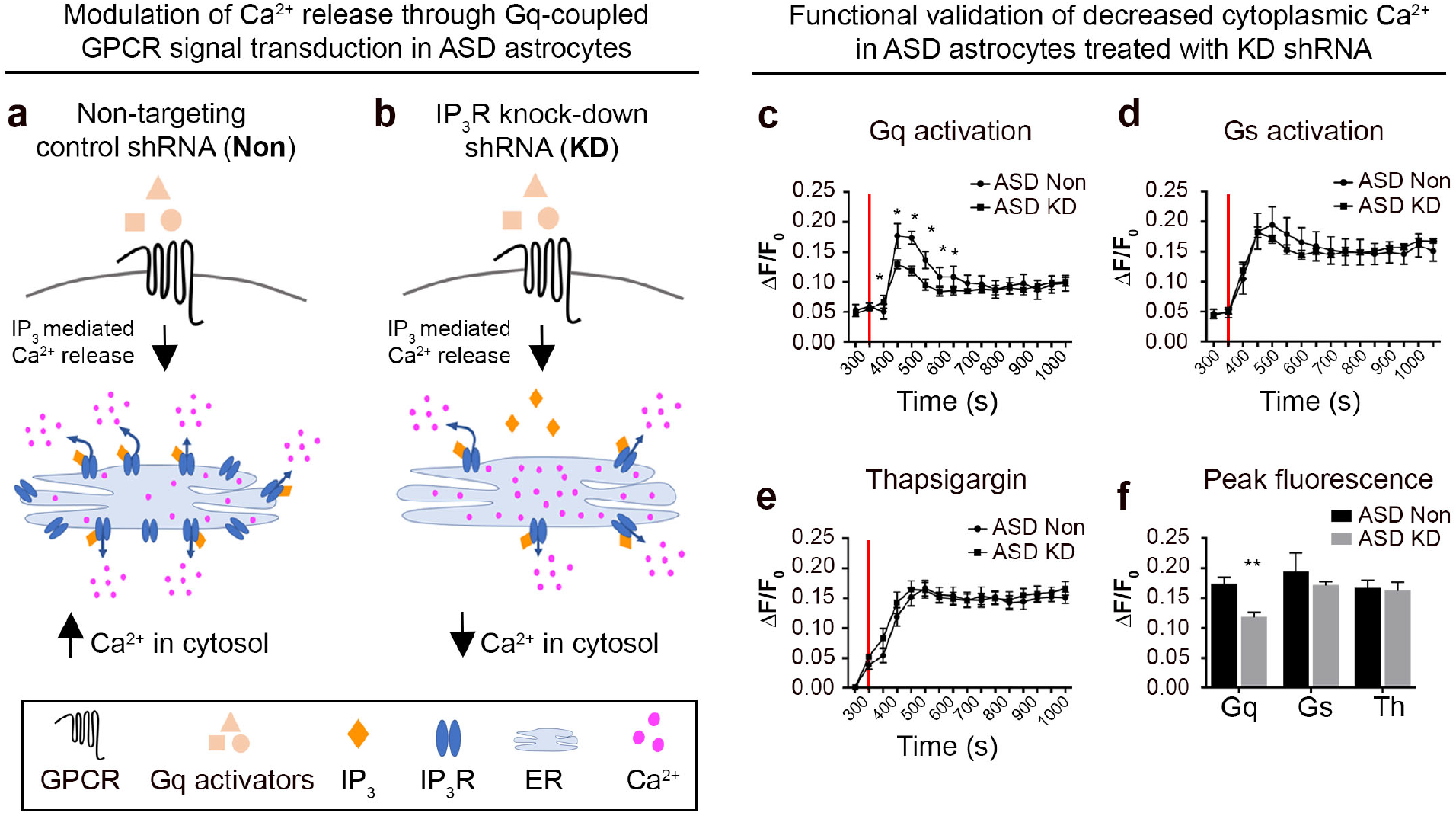
Modulation of Ca^2+^ signaling in ASD astrocytes. **a,b**, To reduce evoked increases in cytosolic Ca^2+^ from internal stores, we transduced ASD astrocytes with an shRNA lentivirus to knockdown IP_3_Rs (KD). As a control, ASD astrocytes were infected with a non-targeting shRNA lentivirus (Non). **c**-**f,** To validate Ca^2+^ modulation by knockdown of IP_3_Rs, we optimized a high throughput Ca^2+^ mobilization assay (see Methods). **c,** ASD KD astrocytes responded to the application of a cocktail of Gq activators (red line, 50 μM DHPG, 50 μM norepinephrine, 50 μM ATP, and 50 nM Endothelin 1) with diminished Ca^2+^ mobilization compared to ASD Non astrocytes. **d,** ASD KD astrocytes did not display reduced Ca^2+^ mobilization in response to activation of Gs GPCR signal transduction (12.5 nM forskolin), which does not rely on IP_3_R mediated Ca^2+^ release from the ER, indicating the specificity of the KD system. **e,** No differences were detected in the total Ca^2+^ concentration stored in the ER between ASD KD or Non astrocytes. Application of thapsigargin (2 μM, red line) depleted the ER Ca^2+^ and inhibited reuptake by Sarco/ER Ca^2+^-ATPase (SERCA) Ca^2+^ pumps. **f,** ASD KD astrocytes showed lower Ca^2+^ responses to Gq activation compared ASD astrocytes, which indicates that IP_3_R reduction in ASD astrocytes modulated evoked increases in cytosolic Ca^2+^ from internal stores without changing its total concentration in the ER (Gq activation vs Thapsigargin).

To validate the reduction of Ca^2+^ mobilization in ASD KD astrocytes, we optimized a high throughput fluorescent screening assay in a 384-well format that enabled application of stimulators under the same conditions with no time lag between wells. CTRL or ASD astrocytes were loaded with a fluorescent indicator dye enhanced for detection in high throughput systems (Fluo-8, Abcam). We found that knockdown of IP_3_R in ASD astrocytes functionally reduced cytosolic Ca^2+^ upon activation of Gq signal transduction compared to ASD astrocytes treated with the non-targeting shRNA lentivirus (**Fig. 6c**). Notably, activation of Gs linked GPCR signal transduction remained intact between ASD KD astrocytes and ASD Non astrocytes, indicating the specificity of our experimental system (**Fig. 6d**). Lastly, IP_3_R shRNA lentivirus did not affect the total Ca^2+^ concentration stored in the ER (**Fig. 6e**) as thapsigargin depleted ER Ca^2+^ stores and inhibited reuptake by Sarco/endoplasmic-reticulum Ca^2+^-ATPase (SERCA) Ca^2+^ pumps. Quantification of peak fluorescence of each group showed that ASD KD astrocytes had lower Ca^2+^ responses (shown with Gq activation) while same Ca^2+^ concentrations (thapsigargin application) compared to ASD Non astrocytes (**Fig. 6f**).

### ASD astrocytes with modulated Ca^2+^ signaling do not induce impaired network activity and memory

Upon functional validation of Ca^2+^ modulation in ASD astrocytes, we tested whether attenuation of Ca^2+^ mobilization prevented dysfunction induced by ASD astrocytes. To address this, we co-cultured E16-18 WT hippocampal neurons with Non or KD shRNA infected ASD astrocytes. Similar to the experiments in **Fig. 5**, ASD Non astrocytes decreased mean firing rate, synchronization, and total number of network bursts in WT hippocampal neurons when compared to CTRL Non astrocytes (**Fig. 7b-e**). Unlike ASD Non astrocytes, ASD KD astrocytes behaved similar to CTRL Non astrocytes in these co-cultures suggesting that attenuation of cytosolic Ca^2+^ levels has a protective effect (see **Supp Video 3**). Representative raw traces of spiking behavior (**Fig. 7a**) and raster plots of network bursts (**Fig. 7d**) visually illustrated that reduced cytosolic Ca^2+^ levels in ASD KD astrocytes had a protective effect on spike and synchronized bursting activity in primary neurons relative to ASD Non astrocytes with high cytosolic Ca^2+^ levels.

**Fig. 7:**
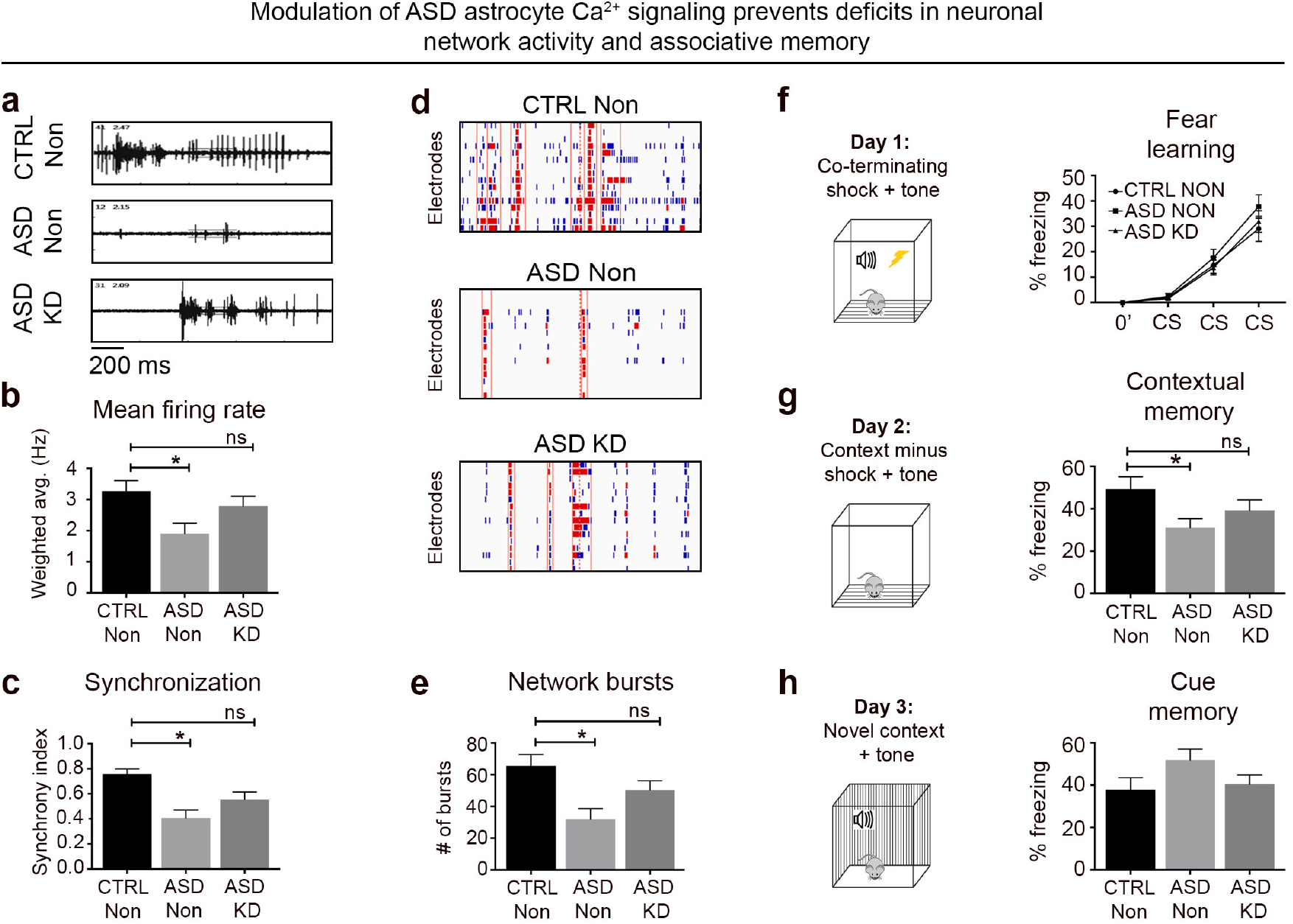
ASD astrocytes with modulated Ca^2+^ signaling do not induce impaired network activity and memory. **a-e**, Human astrocytes were infected with either the non-targeting shRNA lentivirus (Non) or the IP_3_Rs shRNA lentivirus (KD) and co-cultured with primary hippocampal neurons. **a**, Decrease in spiking behavior in untreated ASD Non co-cultures was corrected upon IP_3_Rs KD in ASD astrocytes. **b**, As in Fig. 5, ASD co-cultures displayed decreased mean network firing rate when compared to CTRL co-cultures. Modulation of ASD astrocyte Ca^2+^ mobilization conferred protection against deficits in mean firing rate (See also **Supp Video 3**). **c**, Similarly, ASD Non co-cultures displayed lower synchronicity that was corrected in ASD KD co-cultures. **d**, Knockdown of IP_3_R rescued the phenotypes in network burst number (red hash marks surrounded by red boxes) and spikes per bursts (red hash marks) in the ASD astrocytes. **e**, Unlike untreated ASD astrocytes, ASD KD astrocytes displayed similar number of network bursts when compared to CTRL astrocytes. These findings suggest that fine-tuning ASD astrocyte Ca^2+^ levels protects against deficits induced by untreated ASD astrocytes. **f**-**h**, We transplanted CTRL Non astrocytes, ASD Non astrocytes, and ASD KD astrocytes into the brains of neonatal *Rag2^KO^* mice. **f**, Similar to the previous experiment, there was no significant difference in the rate of acquisition learning between CTRL and ASD mice. **g**, As in Fig. 4, ASD Non chimeric mice showed reduced freezing behavior when exposed to the fear context relative to CTRL Non chimeric mice. However, this difference was eliminated between ASD KD and CTRL Non chimeric mice, indicating that amelioration of cytosolic Ca^2+^ in ASD astrocytes prevents associative memory deficits. **h**, No significant differences detected in freezing behavior between CTRL, ASD, and ASD KD mice during cue presentation in a novel context. Together, these results indicate that exaggerated Ca^2+^ release from internal stores in ASD astrocytes is responsible for neuronal network activity and memory deficits caused by these cells. MEA: CTRL Non co-cultures *n* = 12 wells, 3 distinct lines; ASD Non *n* = 10-16 wells, 4 distinct lines; ASD KD *n* = 15-16 wells, 4 distinct lines. Fear testing: CTRL Non *n* = 24 male and female mice, transplanted with 2 distinct lines, ASD Non *n* = 22 male mice, transplanted with 4 distinct lines, ASD KD *n* = 24 male mice, transplanted with 4 distinct lines. CTRL Non: Control human astrocytes infected with non-targeting shRNA lentivirus ASD Non: ASD patient astrocytes infected with non-targeting shRNA lentivirus ASD KD: ASD patient astrocytes infected with IP_3_R shRNA lentivirus

We next asked if ASD KD astrocytes induced associative memory deficits in chimeric mice similar to ASD astrocyte chimeric mice (**Fig. 4**). To test this, we transplanted neonatal *Rag2^KO^* mice with ASD astrocytes infected with IP_3_R KD shRNA or Non shRNA lentivirus and CTRL astrocytes Non shRNA lentivirus. We compared hippocampal learning and memory in three groups of mice: 1) CTRL Non astrocyte chimeric mice, 2) ASD Non astrocyte chimeric mice, and 3) ASD KD astrocyte chimeric mice. Similar to fear conditioning experiments in **Fig. 4**, all three groups acquired the association between the audible tone (CS) and the mild electrical foot shock (US) at the same rate (**Fig. 7f**). Consistent with the previous result, ASD Non astrocyte chimeric mice displayed deficits in contextual associative memory relative to CTRL Non astrocyte chimeric mice (**Fig. 7g**). The attenuated fear memory phenotype was not present in ASD KD chimeric mice as these mice froze at a similar rate compared to CTRL Non chimeric mice during memory assessment on second day of the test (**Fig. 7g**).

Together, these results suggest that altered Ca^2+^ signaling in ASD astrocytes account for impaired network activity and hippocampal-dependent behavior caused by these cells.

## Discussion

Our study identifies aberrant Ca^2+^ signaling in ASD astrocytes as a mechanism that contributes to specific behavioral and neuronal deficits. Our data provide insight into how astrocyte dysfunction might contribute to behavioral outcomes of ASD, and potentially revolutionize our understanding of disease pathogenesis.

Our data demonstrate that ASD astrocytes induce repetitive behavior as well as deficits in memory and synaptic plasticity, which reflects changes in neuronal network dynamics. To determine disease specific properties inherent to patient derived astrocytes, we isolated astrocytes from iPSC-derived organoids and expanded them in astrocyte selection media (**Fig. 1**). When these human ASD astrocytes were transplanted into postnatal brains, chimeric mice displayed perseverative digging behavior (**Fig. 4g,h**). This chimeric animal model suggests that ASD astrocyte transplantation induces abnormal behavioral phenotypes relevant to specific aspects of autism behavioral pathology in humans. Similarly, ASD astrocyte chimeric mice exhibited attenuated associative memory (**Fig. 4a-d**). In an independent experiment, where neural progenitor cells (NPCs) from other ASD patient lines were transplanted into neonatal brains, we were able to replicate this defective memory phenotype (**Supp Fig. 9-11**). Because the NPC transplantation occurred at a time window when astrocyte generation peaks within the postnatal brain, the vast majority of CTRL and ASD NPCs terminally differentiated into astrocytes in the adult chimeric brains (**Supp Fig. 10**).

Together, this suggests that astrocyte dysfunction causally induce behavioral features in chimeric mice. That said, the heterogeneity of astrocytes isolated from organoids, as well as how distinct subpopulations are affected in ASD, remains to be explored. While our organoid protocol is devoid of serum, astrocytes were maintained in 2% fetal bovine serum for expansion and maturation after dissociation from organoids. Although our data suggests that the cultured astrocytes are not overtly reactive (**Supp Fig. 2**), we cannot exclude the possibility that serum, which is not physiological and only present upon breakdown of blood brain barrier *in vivo*, might select for a certain phenotype or subpopulation.

We also found that ASD astrocyte mice manifest altered LTP relative to age matched human CTRL astrocyte mice (**Fig. 4e,f**). This is consistent with the critical role of astrocytes in proper communication. Astrocyte release of ATP suppresses synaptic transmission and as a result regulates the dynamic range for LTP generation^67^. Likewise, astrocytes have been implicated in formation of new memories through modulating LTP (21), and experimental manipulations to LTP induce behavioral impairments associated with this disease^68, 69^. We, therefore, specifically measured LTP in ASD chimeras. LTP is composed of two different phases^70^: protein synthesis-independent early-phase LTP (E-LTP) and protein synthesis-dependent late-phase LTP (L-LTP). In our study, transplantation of ASD astrocytes disrupted initial phase of L-LTP in hippocampal slices dissociated from ASD astrocyte chimeric mouse brains when compared to control CTRL astrocyte chimeric brain slices. Given that LTP represents the cellular mechanism of memory, these results directly link ASD astrocyte induced circuit disruption with defective associative memory observed in ASD astrocyte chimeric model.

Immune system integrity is suggested to be a player in ASD^71^. In our experiments, *Rag2*^KO^ background is shared between ASD astrocyte and CTRL astrocyte engrafted mice. For chimera studies, use of immune compromised mice as graft hosts is necessary to avoid rejection^44^. Specifically, T-cell and B-cell development is arrested in *Rag2*^KO^ mice. The majority of studies assessing B-cell number and function did not detect any abnormalities in ASD cases^72^. A study found higher number of T-lymphocytes as well as microscopic blebs in the perivascular space in postmortem ASD brains when compared to a healthy control group^73^. These blebs were commonly associated with astrocytes, raising the possibility that there is an active immune response in the ASD brain that targets astrocytes^73^. However, whether this dysregulation is primary or secondary to the disease is not clear. Even if immune system integrity turns out to be a key player in ASD pathogenesis, it surely cannot be the sole factor responsible for all behavioral abnormalities. Our studies provide evidence that ASD astrocytes induce behavioral features in chimeric mice, suggesting a causal contribution of astrocytes to the disease independent of a compensatory response that could also be present in the already diseased brain.

It has been posited that the inability of the ASD brain to properly coordinate or synchronize neuronal activity contributes to behavioral impairments^74, 75^. To provide insight into mechanisms of how plasticity and behavioral phenotypes are induced in ASD astrocyte chimeric mice, we matched the *in vivo* setting by co-culturing ASD astrocytes with mouse hippocampal neurons. These co-culture conditions are the closest to our *in vivo* chimeric models in terms of cellular interactions. Here, we show that ASD astrocytes cause reduced spine density and network activity in hippocampal neurons *in vitro*. Synaptic plasticity takes several forms, including modification of synaptic strength, spine structure and density. Indeed, neuronal connectivity and spine densities are intimately dependent on astrocyte function^14, 76^. Further, ASD postmortem brains exhibit aberrant spine density in frontal, temporal, and parietal lobes^77–79^. Spine density most likely operates in an optimal range to subserve proper cognitive function; thus too many or too few spines is deleterious^80^. Consistent with this, accompanying the phenotype in structural plasticity (reduced spine density in **Fig. 5m-o**), we also found reduced network and synchronous activity in neurons co-cultured with ASD astrocytes compared to those co-cultured with CTRL astrocytes (**Fig. 5a-l**). These experiments have complemented our chimera experiments and support the conclusion that ASD astrocytes induce specific cognitive and behavioral deficits by influencing the structural and functional properties of neurons. Thus, our data suggest a causal contribution of astrocyte dysfunction to ASD through altering neuronal connectivity and plasticity. Our results identify exaggerated evoked Ca^2+^ signaling as an inherent mechanism that contributes to ASD astrocyte mediated phenotypes. We found that proteins that serve in Ca^2+^ signaling were differentially regulated when protein profiling of ASD astrocytes from 9 patients were compared to CTRL astrocytes from 8 subjects. Two-photon Ca^2+^ imaging in transplanted cells *in vivo* (**Fig. 3**) confirmed an aberrant Ca^2+^ mobilization within several distinct patient ASD astrocytes compared to CTRL astrocytes. *In vivo* Ca^2+^imaging of transplanted human astrocytes in awake behaving mice showed that ASD astrocytes exhibited significantly heightened responses to environmental stimuli, as seen both in significantly increased Ca^2+^ elevation as compared to CTRL astrocytes and highly significant increases in all ASD cells in response to the startle stimulus despite decrease or unchanged Ca^2+^ levels. These exaggerated Ca^2+^ activities in ASD astrocytes suggest that Ca^2+^ regulation in these cells may be dysfunctional, leading to aberrant Ca^2+^ activity found in our *in vitro* and *in vivo* experiments and resulting in behavioral abnormalities.

There is an abundance of evidence that supports the theory that mutations along common neurodevelopmental pathways are disrupted in ASD. Recent sequencing studies suggest distinct mutations cluster on neural communication and plasticity networks^81–83^. Consistent with these findings, it was possible to correct deficits in multiple ASD mouse models by targeting common signaling hub critical for plasticity^68, 84^. Further, neurons derived from several nonsyndromic ASD patients and maintained in 2D culture conditions demonstrated defects in Ca^2+^ signaling as well synaptic plasticity^85^. We found that when evoked Ca^2+^ responses were attenuated by knockdown of IP_3_Rs in ASD astrocytes prior to transplantation, no fear memory phenotype was observed in ASD astrocyte chimeric mice (**Fig. 7f-h**). Full genetic knockout of IP_3_Rs eliminates nearly all somatic Ca^2+^ transients, while having no effect on neuronal excitability, synaptic currents, or synaptic plasticity^66, 86^. Thus, modulation of Ca^2+^ signaling with this system reduces non-specific adverse effects to general electrophysiology properties. We also show that attenuation of Ca^2+^ mobilization in ASD astrocytes in co-cultures prevented all neuronal network activity phenotypes previously observed in ASD astrocytes/hippocampal neuron co-cultures (**Fig. 7a-e**, see also **Supp Video 3**). Increases of astrocyte Ca^2+^ evoked by receptor agonists such as glutamate and GABA, or by uncaging of Ca^2+^ or IP_3_, were reported to release gliotransmitters from astrocytes that regulate a wide range of processes in neurons^87, 88^. Increased Ca^2+^ concentration in ASD astrocytes may trigger the release of gliotransmitters, which activates circuits that underlie behavior. Glutamate has been suggested to be one of those gliotransmitters^88, 89^. However, ASD astrocytes did not exhibit significantly increased glutamate releases compared to CTRL astrocytes (**Supp Fig. 12**). While current assays might not detect subtle but physiologically important changes in glutamate release, another possibility is that other gliotransmitters might mediate evoked Ca^2+^ responses of ASD astrocytes.

## Conclusion

Our study identifies aberrant Ca^2+^ signaling in ASD astrocytes as a mechanism that contributes to specific neuronal and behavioral deficits. Our study not only provides mechanistic insight into ASD by showing the causal role of astrocytes to disease pathology but also defines specific physiological roles for astrocyte Ca^2+^ mobilization in associative memory and coordination of network activity. Many cognitive disorders are associated with alterations in neuronal connectivity and synaptic plasticity^90–92^. Mechanisms that regulate these processes are therefore fundamental to understanding higher brain functions. Thus, in addition to providing insights into the mechanisms of ASD, our data also have broad implications for understanding the underlying mechanisms of various other neurodevelopmental diseases.

## Methods

### Materials availability

All unique/stable reagents generated in this study are available from the Lead Contact with a completed Materials Transfer Agreement.

### Data and code availability

The datasets supporting the current study have not been deposited in a public repository because data is embargoed until publication acceptance at which point the dataset will be publicly available from the proteomics databases PRIDE and/or PepditeAtlas, as consistent with editorial standards.

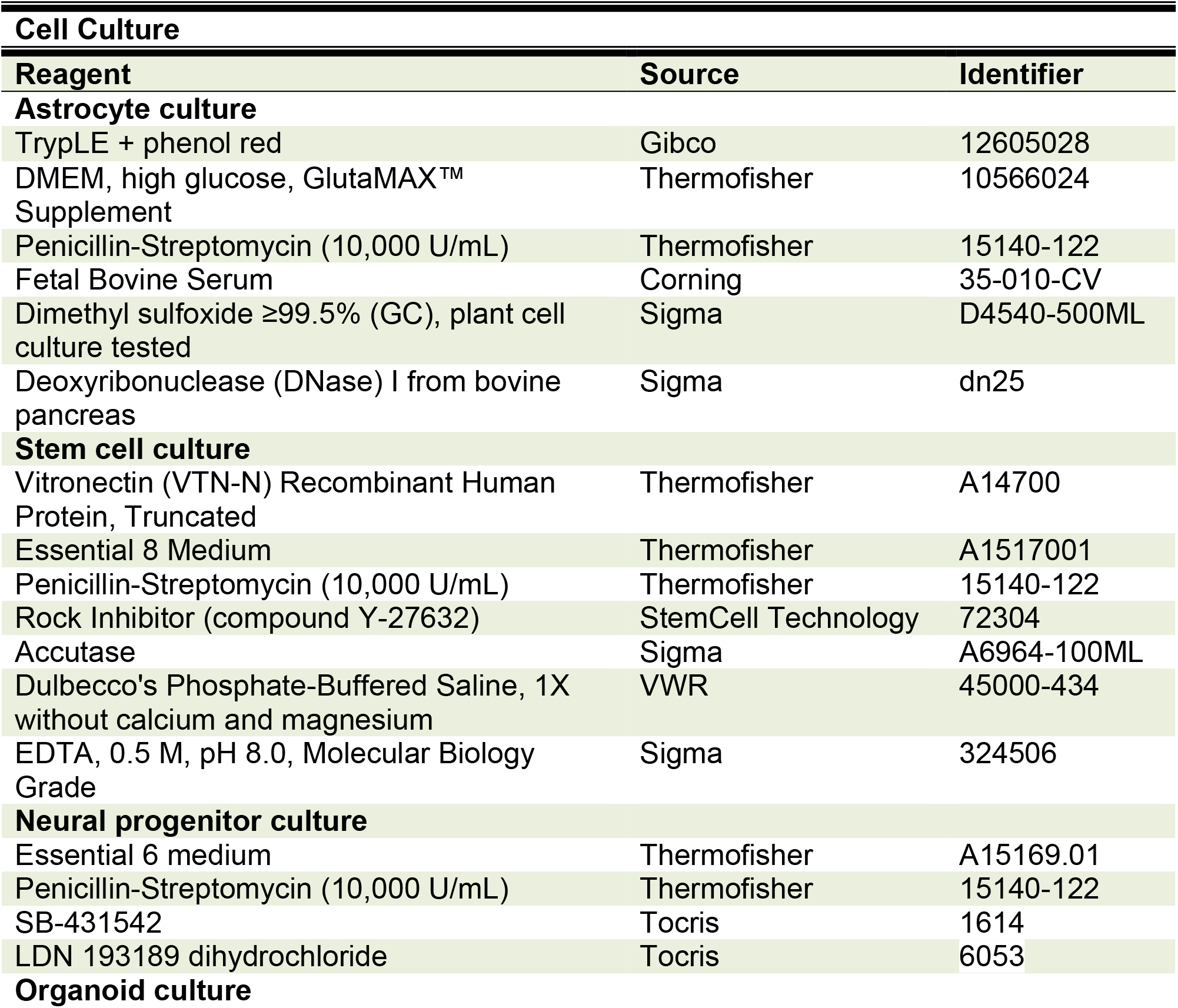
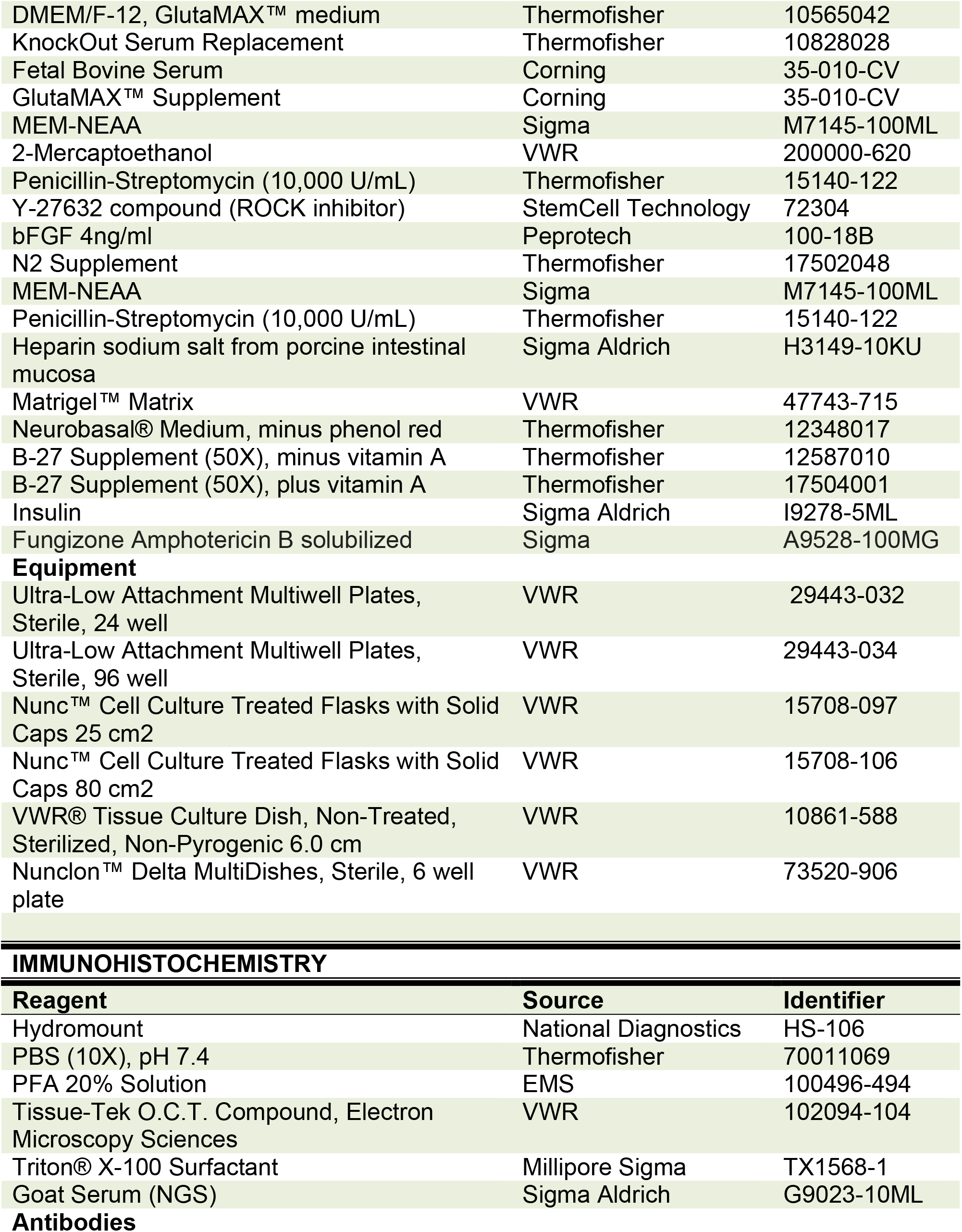
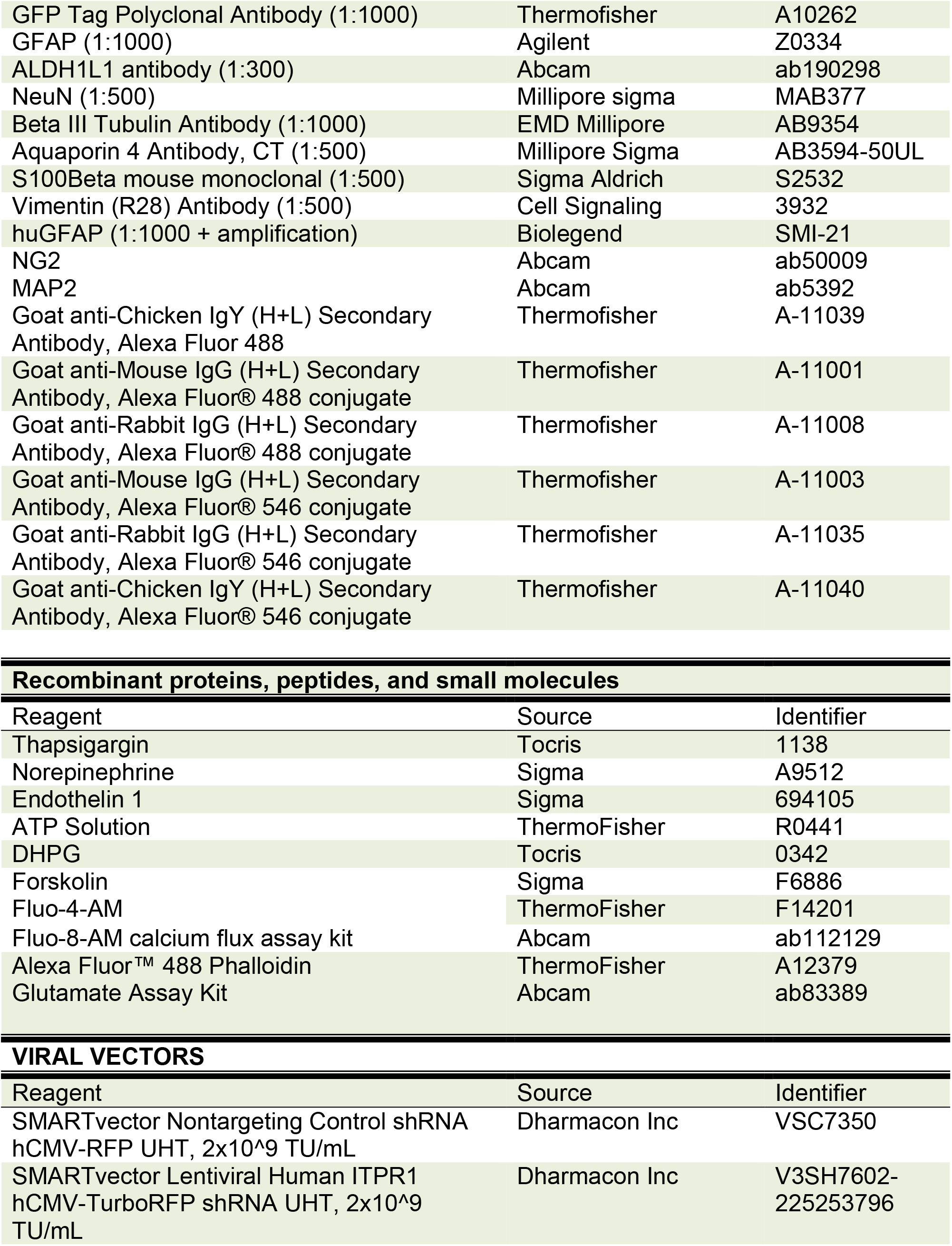
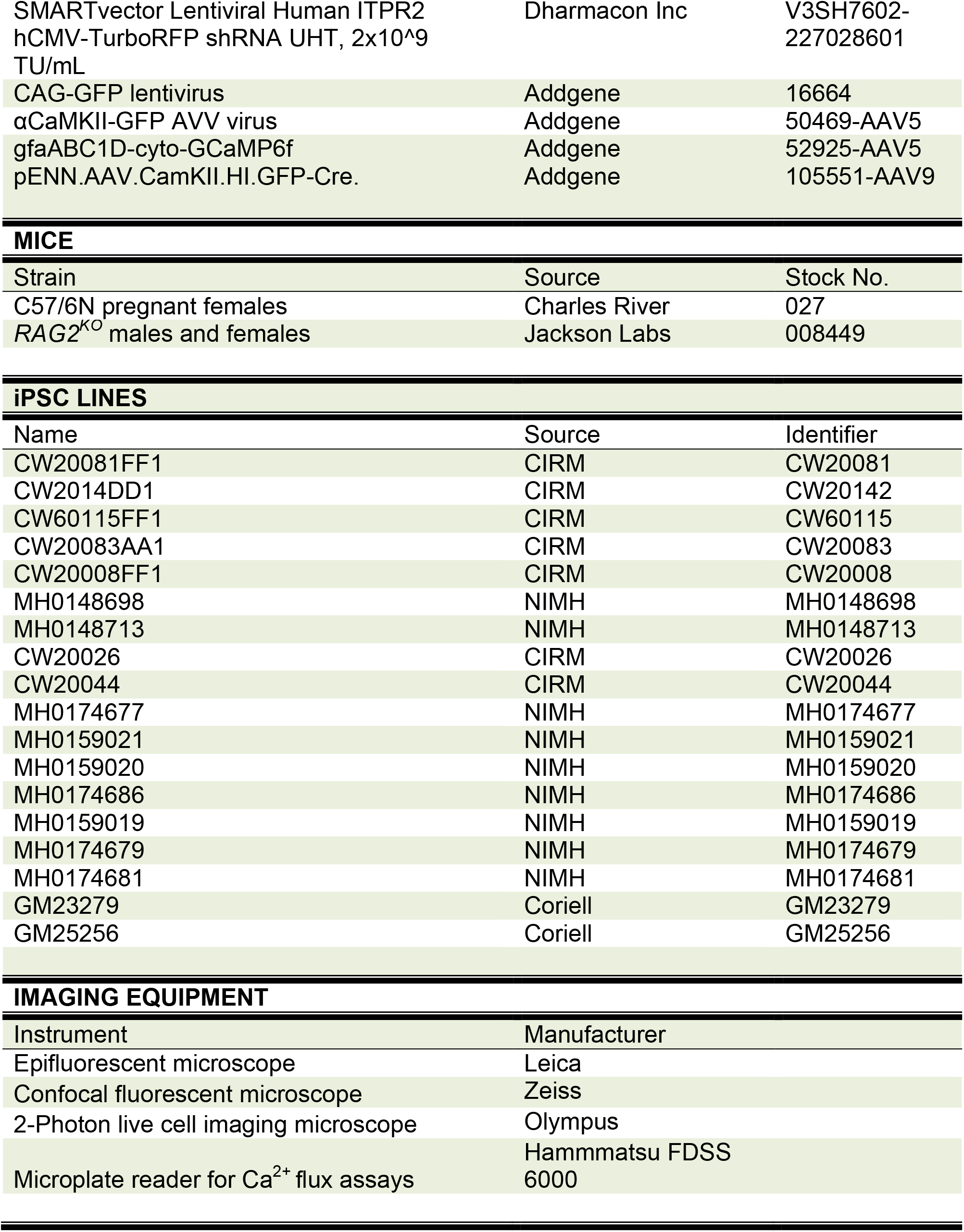
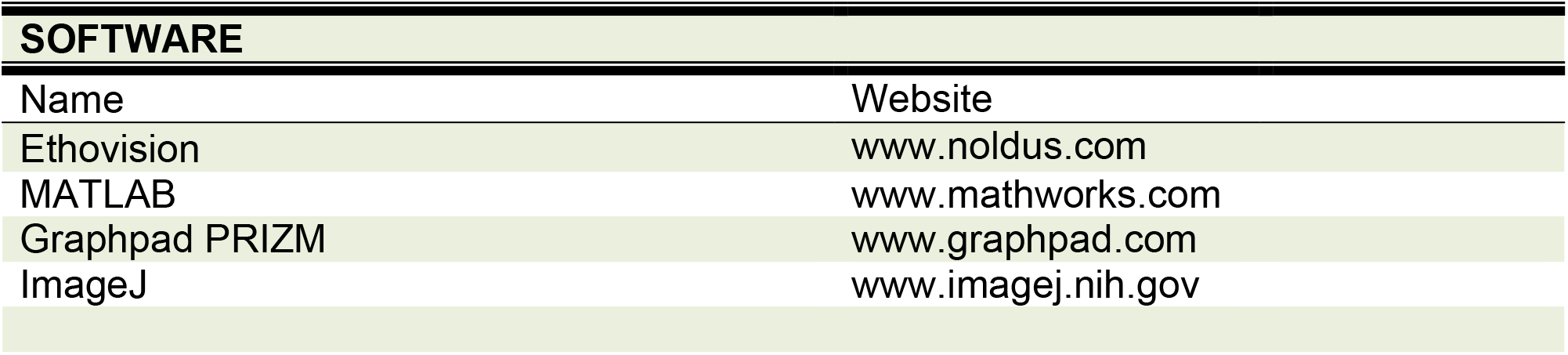
Key resources tables.

### Mice

Animal experiments were conducted in accordance with the ethical guidelines of the National Institutes of Health and approved by the Institutional Animal Care and Use Committee at Weill Cornell Medicine. All animals were group housed with littermates and permitted free access to food and water. Experimental results compare *Rag2^KO^* immunocompromised mice transplanted with cells derived from ASD patients or healthy CTRL subjects. Neuronal cell cultures were dissociated from E16 pregnant C57/6N females purchased from Charles River. See key resources table for more information.

### Induced pluripotent stem cells (iPSCs)

We purchased most iPSC lines from NIH and CIRM repositories. Two CTRL lines were purchased from Coriell Institute. Each repository characterized and validated cells as pluripotent and performed karyotyping to ensure the genomic integrity of each reprogrammed line. ASD lines MH0148698 and MH0148713 from an NIMH collection (Study 116) are genetically characterized and published^93^. Please see Supplementary Tables 1-3 for patient as well as genetic information. Key resources table includes information for purchasing. Supplementary Table 4 provides a list of lines used per experiment.

### Whole Exome Sequencing (WES)

We collected DNA from astrocytes and used NovaSeq6000 to generate 2x151bp paired-end reads at an average depth of 30X. For genetic analysis, we performed whole exome sequencing analysis in all CTRL and ASD lines used here. We screened our detected variants (both SNV/indel and CNV) in our cases and controls that overlapped known variants in ClinVar that have been implicated in autism with a clinical significance of “Pathogenic” or “Likely Pathogenic” resulting in a screening panel of 2367 SNVs/indels and 385 CNVs. We found no relevant hits for SNVs/indels in either our cases or controls with these established autism variants. We detected 25 CNVs in our cases and 31 CNVs in controls that had a 50% reciprocal overlap with ClinVar CNVs, all of which were duplications (aside from one deletion) that overlapped non-critical regions of “Pathogenic” or “Likely Pathogenic” CNVs. Moreover, we compiled a screening panel on the gene level comprised of 322 autism genes obtained from SFARI Gene database (from the high confidence category) as well as human genes from the MGI database that have been linked with autism. Across these 322 genes, we identified 33 rare and potentially impactful variants in our case samples and 37 variants in our control samples. All of these variants, with the exception of 2 in-frame insertions, are missense variants with a ‘moderate’ predicted impact. None of these variants harbor a “Pathogenic” or “Likely Pathogenic” classification by ClinVar at the variant level. Supplementary table 2 displays the counts of hits for LGD SNVs and LGD CNVs. Supplementary table 3 displays variants in cases and controls and which individual carries them.

### iPSC cultures

All iPSC lines were thawed and directly plated on vitronectin (1µg/cm^2^) coated 6-well tissue culture treated plates. ROCK inhibitor (10 µM) was added to culture media for the first 24 h after thawing. Otherwise, iPSC cultures were maintained in E8 base media plus E8 supplement, and 1X penicillin/streptomycin antiobiotic and incubated at 37°C with 5% CO_2_ humidified air. Media changes occurred 6 days a week. We surveyed wells daily and passaged with 0.5 mM EDTA once colonies reached 70-80% confluency. Optimal iPSC lines formed spherical colonies with slightly raised edges. If spontaneous and excessive differentiation occurred, we manually isolated undifferentiated colonies under a dissection scope with sterile pipette tips and transferred to newly vitronectin coated 6- or 12-well culture treated plates. No ROCK inhibitor was added when manually transferred.

### Neural progenitor expansion

iPSC colonies were enzymatically dissociated with 1X Accutase^®^ solution in dulbecco′s phosphate-buffered saline (DPBS) without Ca^2+^ or Mg^2^ at 37°C for 10 min. Next, cells were transferred to a 15-mL falcon tube, diluted in E8 media to inactivate the enzymatic reaction, and centrifuged @1000 RPM for all lines. iPSCs were resuspended in E8 media plus ROCK inhibitor and plated in 6-well plates that were precoated with 1:40 matrigel at a density of 1.5x10^6^ cells per well. Once wells reached ∼70% confluency, media was changed from E8 to E6 to help promote the formation of a uniform neuroepithelial layer. In adherence to the dual SMAD inhibition protocol^94^, E6 media was supplemented with SB-431542 and LDN 193189 factors. Media changes occurred daily and cells were observed for presence of a neuroepithelial sheet, in which cell bodies became smaller and more densely packed. This typically occurred in our hands between 5-8 days in E6+factor media. Timing of neuroepithelial development varied slightly between each independent line. Between days 5-8, progenitors were enzymatically dissociated with Accutase^®^ solution, prepared as a single cell suspension in DPBS, and housed on ice prior to transplantation into the brains of *Rag2^KO^* newborn pups between postnatal day 1 and 3 (P1-3).

### Organoid development

To obtain ASD patient astrocytes without biasing differentiation, we integrated two distinct strategies for successful generation of human iPSC-derived astrocytes (29, 30). 3D cortical spheroids spontaneously generate astrocytes, which express similar protein profiles as purified human brain astrocytes^29, 31^. However, by design, directed protocols like the cortical spheroid method, call for factors that potentially prejudice differentiation. Here, we adapted an undirected-differentiation organoid system^32, 33^. We cultured iPSCs into three-dimensional embryoid bodies, induced neural fate specification, and allowed free floating tissue to expand for 75 days in 10 cm sterile tissue culture dishes. During this expansion phase, organoids were maintained in media (DMEM-F12, Neurobasal medium, N2 supplement, B27 supplement + vitamin A, insulin, GlutaMAX supplement, 2-mercaptoethanol, MEM-NEAA, penicillin-streptomycin, and fungizone) and agitated at 60 RPM on an orbital shaker to ensure nutrient diffusion. Because this organoid protocol does not use excessive patterning factors or small molecules, it likely preserves disease-specific signatures unique to patient astrocytes. Cell fate specification is temporally regulated in cortical development and is defined by the sequential appearance of neurons followed by glia^95^. Undirected organoid protocols recapitulate the temporal sequence of cortical development seen in developing embryos^34^, and so, patient organoids spontaneously generate astrocytes in an environment that mimics the early ASD brain. Specifically, ASD astrocytes develop in the presence of genetically matched ASD neurons. This is important because astrocytes require early interaction with neurons to induce expression of critical receptors and trigger temporally regulated activation patterns^35^.

### Astrocyte expansion

To expand CTRL and ASD astrocytes in 2D culture, we enzymatically dissociated organoids at 75 days *in vitro* (DIV). We next selected for astrocyte enrichment with cell culture medium supplemented with glucose and low serum content (2%) as previously described^30^. Transcriptomic profiles of iPSC-derived astrocytes expanded with this method resembled that of human fetal astrocytes. In fact, 90% of the enriched genes were shared between iPSC-derived astrocytes maintained in low serum and human fetal astrocytes^30^. Organoids were enzymatically dissociated in a 0.1% TRYP-LE solution for 15 min and washed 2X with astrocyte selection media (DMEM supplemented with 4.5 g/L of D-glucose, 2mM GlutaMAX™, 2% fetal bovine serum and antibiotic and fungicide). Gentle trituration with a fire polished glass pipette was used to obtain a single cell suspension. This suspension was plated on tissue culture treated 6-well plates at a density of 1.5 million cells per well. Astrocyte selection media supplemented with 10 µM ROCK inhibitor was added to the cells the first 24 h after dissociation. Astrocyte expansion occurred in 2D adherent culture conditions with D-glucose, GlutaMAX™, 2% fetal bovine serum and antibiotic and fungicide. Previous studies have shown that high levels of serum (>10%) result in reactive phenotypes that are not representative of astrocytes *in vivo*^96, 97^. In our hands, minimal serum exposure enhanced astrocyte survival and proliferation and permitted large-scale astrocyte production without causing reactivity. Neurons co-cultured with CTRL astrocytes derived in this way survived and retained neuronal network activity that was similar to neurons without human astrocytes (refer to **Fig. 5**). These astrocytes were enzymatically dissociated with TRYP-LE solution, prepared as a single cell suspension in DPBS, and housed on ice prior to transplantation into the brains of *Rag2^KO^* newborn pups between postnatal day 1 and 3 (P1-3).

### Astrocyte/neuron co-cultures

For the preparation of primary neurons, cells were dissociated from the brains of E16 C57/6N embryonic mouse pups. After the removal of adherent meninges, hippocampi were removed and dissociated into a single cell suspension with trituration following incubation for 15 min with 0.1% TRYP-LE as outlined in^98^. Cells were counted with a hemocytometer and a mixture of neurons was added to a mixture containing astrocytes in a 4:1 ratio. This mixed cell suspension was plated onto 24x50mm, No. 1.5 sterilized glass coverslips or 48-well multielectrode array culture plates that were precoated with 0.01% poly-D-lysine. Cultures were maintained for the first 24 h in 50% neural basal media supplemented with B27 and 50% astrocyte base media. After the initial plating, subsequent feeds contained 100% neural basal media supplemented with B27, 1% glutamine, and penicillin/streptomycin antibiotic. Co-cultures were housed at 37°C in a humidified 5% CO_2_ containing atmosphere.

### Transplantations

Astrocyte or neural progenitor human chimeras were prepared as described^42, 46^ with minor modifications. Prior to transplantations, cells were resuspended in the DPBS at a concentration of 5x10^5^ cells/µl and maintained on ice. Newborn *Rag2^KO^* pups (P1-3) were cryoanesthetized and 2.5x10^5^ human derived cells were injected in equal amounts (0.5 µL) into the forebrain at 4 locations. Bilateral injections were made immediately anterior and posterior to bregma. Injections were made by hand using a Hamilton syringe (30 gauge) and inserted gently through the skin and skull. Pups were monitored 2X a week until weaning and then separated by gender.

### Behavioral testing

Aged matched cohorts were housed in temperature-controlled rooms with a 12-h light/dark cycle. Cages were changed weekly; care was taken not to test animals on the same day as cage change. To further minimize confounding results due to stress, mice were handled for 3 consecutive days prior to the start of testing. When possible, the experimenter was blinded to engraftment group. For all behavioral assays, mice were habituated to the testing room for a minimum of 30 min, and testing was performed at the same time of day except where noted. Animals were 3-5 months of age at the time of testing.

#### Open Field

We tested general ambulatory activity and exploratory behavior of chimeric mice with an open field test (San Diego Instruments). Four boxes, each measuring 50cm × 50cm, allowed for the simultaneous testing of up to four mice. Each box contained a 16 × 16 photobeam configuration that reported beam breaks in x, y, and z planes. The associated PAS software accurately recorded all beam interruptions in real time to report central and peripheral localized activity, ambulatory movements, fine movements, and rearing.

#### Sociability

To test sociability in mice, we used the 3 chambered social approach assay with slight modifications^15^. Testing occurred in a 3-chambered apparatus where animals were free to access to all chambers during testing. The test was divided into two phases: habituation and social preference. During the habituation phase test animals were placed in center of apparatus box that contained two identical wire mesh cylinders at each end chamber. Animals were monitored for 10 min during each phase. Next, a stimulus mouse was placed in one of the wire mesh cylinders. Because testing occurred over multiple days, placement of the social cylinder was counterbalanced throughout testing to eliminate any bias toward any chamber in the apparatus. The time spent in the social or non-social chamber was recorded over a 10-min testing period as well as the time spent interacting with the cylinder that contained the social stimulus. Automated analysis was performed with Noldus Ethovision XT software.

#### Associative learning and memory

Context and cue dependent fear conditioning is an associative learning task where mice learn to associate a neutral conditioned stimulus (CS; audible tone, 70 dB, 30 sec duration) with an aversive unconditioned stimulus (US; mild electrical foot shock, 0.7 mA, 1 sec duration) and display a conditioned response (CR; freezing behavior)^99^. The CR or freezing behavior can be used as an index of learning and memory. After repeated pairings of CS and US, the animal learns to fear both the tone (cue-dependent memory) and training context (context dependent memory). Chimeric mice learned to associate a neutral conditioned stimulus (CS; audible tone) with an aversive unconditioned stimulus (US; mild electrical foot shock) during a 5-min training session. On day 1, mice were permitted a 2-min habituation period followed by 3 successive 30 sec trials of coterminating shock and tone. On day 2, we assessed each group’s ability to recall the association between the aversive stimulus and the training box (context-dependent associative memory). Chimeric mice were placed in the training boxes for 5 min. The first 2 min of freezing were reported. On day 3, we assessed each group’s ability of recall the association by placing animals in a novel context and administering only the audible tone. We reported average freezing % during the first tone presentation. On all testing days, % freezing behavior was a readout for associative memory. Freezing percentages were determined by automated software from Med Associates Inc.

#### Spatial learning and memory

Hippocampal learning and memory deficits were evaluated using the Morris water maze paradigm as previously described^15^, with additional modifications. Specifically, the water maze apparatus consisted of a 4ft diameter pool filled with opaque water (25°C; colored with non-toxic Crayola® white paint). Visual cues were placed on the walls surrounding the pool and a platform (4 inches in diameter) was hidden below the surface of the water (1 cm). Initial training consisted of four trials per day over four days. Mice were introduced into the pool at variable entry points, with every entry point used over the course of the day. The location of the platform remained constant throughout training period. The mice were given 60 sec to locate platform. On the fifth day of testing, a probe trial was conducted in which the platform was removed over one 60-second trial. Tracking software (Noldus Ethovision XT) was used to record swim speed, total distance traveled, distance traveled in platform quadrant, time spent in platform quadrant, and platform crossings. *<u>Repetitive behaviors</u>*. To test mice for repetitive behaviors we used marble burying. For marble burying, we quantified the number of marbles a mouse buried over a 30-min period in a cage that contained extra bedding at a depth of 5 cm and 28 marbles arranged in a 4  ×  7 grid^58^. Marbles were scored as buried if at least 60% covered.

### Long-term potentiation (LTP)

All electrophysiology experiments were performed on transverse hippocampal slices (400 μm) from 4 to 6-month-old mice. Slice preparation, and ACSF composition were performed as previously described^100^. L-LTP was induced with three trains of 1 s, 100 Hz high frequency stimulation (HFS) with an intertrain interval of 60 s. Bipolar stimulating electrodes were placed at the border of area CA3 and area CA1 along the Schaffer-Collateral pathway. ACSF-filled glass recording electrodes (1–3 MΩ) were placed in stratum radiatum of area CA1. fEPSPs were amplified (A-M Systems Model 1800) and digitized (Digidata 1440; Molecular Devices) prior to analysis (pClamp; Molecular Devices). The initial slopes of the fEPSPs from averaged traces were normalized to those recorded during baseline.

### Immunocytochemistry

Astrocytes were plated on uncoated sterile coverslips at 5x10^4^ cells per well of a 24-well plate. For co-cultures, 7.5x10^4^ neurons were mixed with 2.5x10^4^ astrocytes and plated onto poly-D-lysine coated sterile coverslips. Cells were fixed in 4% PFA for 10 min immediately followed by permeabilization with 0.5% Triton X for 10 min and blocked with 10% Goat Serum. Primary antibodies were incubated overnight at 4°C followed by a 2 h incubation with secondary antibodies at RT. To visualize the nucleus of individual cells, Hoescht was applied in the final wash prior to mounting. *<u>Free floating immunohistochemistry</u>* Cryosectioned free floating sagittal brain sections (30-40 μm thickness) of perfused and fixed mouse brains were collected in PBS and permeabilized in 0.5% Triton X for 10 min, and blocked with 10% Goat Serum. Primary antibodies were incubated overnight at 4°C with slight rocking. The following day, secondary antibodies were added for 2 h at RT. To visualize the nucleus of individual cells, Hoescht was applied in the final wash prior to mounting. In addition to these steps, amplification kits were used for huGFAP immunostaining only.

#### Quantitative immunostaining

ImageJ was used to semi automate the quantification of the colocalization of GFP with ALDH1L1 and GFAP in cryosectioned free floating sagittal brain sections. Composite images of the somatosensory cortex and hippocampus were acquired on epifluorescent microscope (Leica). To process, composite images were split into individual channels and transformed into 8 bit images. Standardized thresholding and processing parameters were applied to all images and then cells were manually counted with the help of the ROI manager tool.

### Spine density

We dissociated primary hippocampal neurons from the hippocampi of C57BL/6N embryos at E16 as previously described with some modifications^98^ and plated 1x10^5^ neurons with 4x10^4^ astrocytes isolated from CTRL or ASD patient organoids on 0.01% poly-D-lysine coated 12 mm round glass coverslips. To visualize dendrites and dendritic spines, we infected neurons with an adeno-associated virus that expressed GFP under the control of the α*CaMKII* promoter. This promoter-specific expression of GFP restricted labeling to only excitatory neurons and their spines. To perform spine analysis, we fixed neurons at DIV18 with 4% PFA, amplified the GFP signal with antibodies and fluorescent immunochemistry. We acquired images by generating maximum intensity projections from z-stacks using optical sections (Olympus confocal microscopy, 0.35 µm per section). To quantify, we identified a 10 μm dendritic segment at least 30 μm away from the soma and counted individual spines using ImageJ.

### Stereology

Stereo Investigator Stereological Software (MBF Bioscience, Williston, VT) was used to estimate the total number of GFP+ human astrocytes in hemispheres of chimeric mice. Chimeric mice underwent transcardial perfusion with 4% PFA at P60 and brains were cryosectioned along the sagittal plane (30μm thick). Every 4^th^ slice was taken for analysis. Slices underwent immunostaining to enhance GFP signal. Sections were traced under low magnification (4X) and counted under high magnification (20X) using the optical fractionator method. Briefly, the software would systematically and randomly choose counting frames within the brain section. Then, it would draw an 80μm × 80μm square around the counting frame. Cells were counted if they were found within that box and only if they did not come into contact with two borders of square frame.

Upon completion of counting, we applied the numerical formula supplied by MBF bioscience to extrapolate the total cell number within each brain hemisphere. To calculate estimates, we multiplied the total number of cells counted by the total volume of the brain slice (mm^2^) divided by the volume of counting square (mm^2^).

### Cranial window implant surgery, recovery, and habituation

Adult (P60+) mice with transplanted human astrocytes expressing GCaMP6f were anesthetized with isoflurane (4% for induction, 1–1.5% vol/vol for maintenance) and placed in a stereotaxic frame (Kopf), with body temperature kept at ∼37 °C with a feedback-controlled heating pad (Harvard Apparatus). After removing the scalp and clearing the skull of connective tissues, a custom-made titanium head-bar was fixed onto the skull with cyanoacrylate adhesive (Krazy Glue) and covered with black dental cement (Ortho-Jet). A circular craniotomy (3-mm diameter) was then performed above the border of primary somatosensory and motor cortices on the right hemisphere (right S1/M1, centered at 2.0 mm lateral from bregma). The bone was removed without damaging the underlying dura. Then a glass cranial window consisting of a 3-mm diameter #1 coverslip (150 µm thickness, Warner Instruments) was place on top of the dura, flush with the skull surface, and sealed in place using tissue adhesive (Vetbond, 3M). The exposed skull surrounding the cranial window was then completely covered with cyanoacrylate adhesive and then with black dental cement to build a small chamber, intended to hold liquid for imaging with water-immersion objective. Extreme care was taken to ensure that the dura experienced no damage or major bleeding before and after cranial window implantation. Mice with damaged dura or unclear window were discarded and not used for imaging experiments. After surgery, animals were returned to their home cages for at least 3 weeks for recovery, to provide ample time for astrocytes to return to their nonreactive state. After one week in recovery, animals were subjected to periodic handling and head-fixation for habituation to the imaging conditions. Animals were housed in reversed light-dark cycle (light on from 7pm-7am), and all habituation, imaging, and behavioral experiments were conducted during the dark/active phase for the animals.

### Two-photon *in vivo* Ca^2+^ imaging

We dissociated ASD or CTRL organoids, expanded the astrocytes in culture, and virally transfected the astrocytes with AAV carrying the genetically encoded calcium indicator GCaMP6f (AAV2/5-*GfaABC_1_D-GCaMP6f*). We engrafted the GCaMP6f-expressing human astrocytes into newborn *Rag2*^KO^ by injecting the cells into the frontal cortex. We raised the transplanted mice to adulthood (P60+) before subjecting them to *in vivo* experiments. We implanted a 3-mm-diameter glass cranial window over the primary somatosensory and motor cortices (S1/M1), attached a titanium head-bar to the skull for head-fixation, and secured the implants with (black) dental cement. We allowed the mice to recover post-surgery for at least 3 weeks to provide ample time for astrocytes to return to their non-reactive state. During this time, we periodically subjected the mice to handling and head-fixation to habituate them to experimental conditions. Finally, we performed *in vivo* two-photon imaging to record Ca^2+^ activity in transplanted human astrocytes within live mice actively responding to the environment. Two-photon calcium imaging was performed with an Olympus multiphoton laser-scanning microscope (FVMPE-RS) with a Ti-Sapphire laser (Spectra-Physics Mai Tai DeepSee) tuned to 920 nm, through a 25X, 1.0 NA water-immersion objective (Olympus XLPLN25XSVMP2). Photograph in **Fig. 3a** shows labeled components of our custom-designed floating platform, developed to provide a tactile virtual-reality environment for head-fixed mice and to enable imaging of actively locomoting animals with minimal motion confounds. The photo shows a mouse situated at the center under a microscope objective and on top of a platform comprised of a 15x15 cm plastic container with 1cm depth, enclosed by a thin sheet of flexible plastic film material as walls, and filled with bedding material from the animal’s home cage. The platform floats on water inside a larger plastic container (40 x 40 x 8 cm). The animal’s head is immobilized by fixation of its attached head-bar to a custom-built holder consisting of a heavy stable base assembled with goniometers and multi-axis micro-positioning devices for making fine adjustments to the position and angle of the animal’s head. While head-fixed, the animal’s body movement on the floating platform directly translates into platform movement, akin to moving on a treadmill or tactile virtual-reality. The fluidity of water allows the forces generated by the animal movement on the platform to be absorbed by the water instead of causing vibration of the head relative to the objective lens and resulting in image motion artifacts. The imaging setup includes an air puff delivery tube (3mm inner-diameter PVC tubing) connected to an air compressor on one end and positioned at 1cm away from the animal’s face/nostril on the other for delivering the startle stimulus. The setup also includes an infrared-sensitive camera and an infrared light for monitoring animal behavior and locomotion in the dark during 2-photon imaging. *<u>Imaging setup</u>* (***Fig. 3c***): (Left upper panel): a representative image of GCaMP6f-expressing human astrocytes in the cortex of a mouse engrafted with cells derived from a CTRL human subject. *Right, upper panel (orange box)*: Ca^2+^ transient recorded from the cell marked by the orange circle on the left image, showing an increase in Ca^2+^ level after the startle/air-puff stimulus (*orange arrow*). Ca^2+^ level was sustained at an elevated level (*gray line/arrow* = +5.5% ΔF/F) for the duration of the recording. Note the images were acquired at 15Hz (15 frames per second) using one-way resonant scanning, at a much higher rate than typical astrocyte two-photon imaging using galvanometer-based scanning (less than 1Hz). This higher acquisition rate has enabled us to record Ca^2+^ fluctuation dynamics in much higher temporal resolution, in addition to allowing for more robust image motion correction by minimizing in-frame motion artifacts. (Left bottom panel): The objective is placed at the end of an articulating arm (representing the Olympus inner-focus articulating nosepiece); it is angled (15 degrees to the vertical) to allow the animal to assume a more natural head position, for ease of locomotion and stress minimization. The video camera icon on the left represents the infrared-sensitive camera recording the animal’s behavior and locomotion (platform position). Videos were acquired at 15Hz to produce one-to-one corresponding frames between the two-photon images and video recordings. (Right bottom panel-gray box): Representative trace of recorded locomotion during imaging, produced by analyzing the infrared video recording and plotting the light intensity change of a select ROI (region of interest) on the floating platform. As animal movement directly translated into platform displacement, any body motion generated by the animal could be tracked with accuracy and high temporal resolution, even heavy breathing could be seen indicated by the brief ticks at the latter part of the recording.

### Data processing and analysis of *in vivo* Ca^2+^ imaging

For the present study, extra care was taken to minimize potential motion confounds, with extensive habituation to reduce animals’ stress and the implementation of a newly designed floating platform that allowed the force of the animal’s movements to be absorbed by the fluidity and buoyancy of the water supporting the platform, especially in the up-down direction to mitigate the hard-to-correct z-axis motions. Resonant imaging at 15Hz also contributed to avoiding in-frame motion confounds and allowing minor between-frame shifts to be effectively corrected by accurate rigid-body transformations using the TurboReg plugin for ImageJ. Motion-registered data were then processed and analyzed following the steps outlined in Srinivasan and Huang *et al*.^48^ An ROI-based approach is used to segment astrocytes in a semi-automated manner based on maximum-intensity projections of background-subtracted image stacks. As compared to previous studies with mouse astrocytes, the transplanted human astrocytes displayed high basal fluorescence levels and tend to be clustered instead of being spread out in typical non-overlapping manner. These factors, combined with the feature of human astrocytes possessing fine processes that are radially distributed around the soma, made it difficult to resolve the fluorescence from processes vs. the soma to distinguish these different compartments. The ROI selected were assumed to encompass both soma and microdomains in processes. The post-startle percent changes in calcium levels were calculated by subtracting the mean fluorescence value from a pre-startle stable period (first 100 timepoints, 6.6 s) from the mean value form a post-startle stable period (last 100 timepoints, 6.6 s), divided by the initial mean F. The cells showing percent increase above +2% were categorized into the Increased Response group, those below -2% into the Decreased Response group, and those in between +2 and -2% in the Unchanged group. The percent change in variance were calculated similarly as above but using the standard deviations as the measure. Statistical comparisons were made using unpaired two-tailed Student’s t test.

### Neuronal network activity

Recordings on multielectrode arrays (MEAs) began at DIV 14. The MEA system (Maestro system from Axion BioSystems) provided a non-invasive extracellular recording method that did not damage neuronal viability *in vitro.* Using the 48-well multielectrode array (MEA) plates from Axion Biosystems, we plated primary mouse neurons with astrocytes derived from 4 ASD patients and 3 control subjects in triplicate. A non co-culture control was added for each MEA recording. This included neurons dissociated from the same litter but without any human astrocytes. These non-human control cultures were subjected to the same conditions as mouse neuron/human astrocyte co-cultures. Each well was precoated with 0.01% poly-D-lysine and seeded with 7.5x10^5^ neurons and 2.5x10^5^ astrocytes. Co-cultures were fed twice a week and measurements were taken at DIV14. Recordings were performed with the Maestro Pro MEA system and spike sorting and analyses performed in AxIS software from Axion Biosystems. Each well contained 16 electrodes that detected changes in extracellular field potentials reflecting the spiking activity of neurons. Electrodes with an average of ≥5 spikes/min were defined as active. Spikes were detected when they were ≥5.5x the standard deviation of the raw signal per electrode. Each plate equilibrated for a minimum of 10  min prior to recording in Maestro Pro Instrument. Recordings occurred within a chamber heated to 37°C with 5% CO_2_.

### High throughput Ca^2+^ detection assay

Astrocytes were seeded on clear bottom black 96-well plates at 1x10^4^ astrocytes per well 24 h prior to recording. The following day, cells were loaded with Ca^2^ indicator dye optimized for use in high throughput systems (1 μM Fluo-8-AM). Astrocytes were incubated with Fluo-8-AM for 30 min at 37°C. The plate was placed in a preheated plate reader (Hammatsu FDSS 6000) and recording occurred in Ca^2^ free HBSS at 37°C. Baseline fluorescence was recorded for 5 min (470–495  nm excitation and 515–575  nm emission). Next, a robotic arm applied G_q_ cocktail, forskolin, and thapsigargin to predetermined wells. The recipe and final concentration of G_q_ cocktail was as follows: 50 μM DHPG, 50 μM norepinephrine, 50 μM ATP, and 50 nM Endothelin diluted in HBSS. Forskokin (final concentration 12.5 nM) and thapsigargin (final concentration: 2 μm) were also diluted in HBSS. After application of stimulators, recording continued for 10 additional min. Recordings were performed in duplicate or triplicate. The peak change in fluorescence amplitude (ΔF) in each well was normalized to the basal fluorescence of that well before stimulation (F_0_). All data are represented as mean of ΔF/F_0_ from each corresponding cell line from the same plate.

### Glutamate Assay

Astrocytes were grown in T-25 flasks with a starting density of 3x10^5^. Supernatants were collected at 96 hours (no media change). Astrocytes were dissociated and counted after media collection for normalization. Glutamate assay was performed according to kit instructions (Abcam, ab83389) and read in a 450-nm microplate reader.

### Proteomics

To determine differences in protein expression profiles of CTRL and ASD astrocytes, we used tandem mass tag (TMT) liquid chromatography mass spectrometry at the Proteomics and Metabolomics core facility of Weill Cornell Medicine. Briefly, proteins were extracted from cell pellets and quantified with the Bradford assay. Samples were enzymatically digested, labeled with tandem mass tags (TMT), and then combined. The combined TMT-labeled peptides were desalted and subject to LC/MS analysis. MS raw data was searched against Uniprot human database using MaxQuant and Perseus Student’s T-test was performed and p values were adjusted with Benjamini-Hochberg (BH) Correction to avoid false positives. All statistical and bioinformatics analyses were performed using the freely available software Perseus (as part of the MaxQuant environment), the R framework, or EdgeR framework. Proteins identified only by site modification or found in the decoy reverse database were not considered for data analysis. The intensities of proteins were used calculate the ratio and the p-value using EdgeR. Differentially expressed proteins were identified as follows: ratio ±1.5(log2), p-value <0.05. Pathway enrichment analysis for categorical data was performed based on a Fisher’s exact test with a Benjamini–Hochberg FDR threshold of 0.02. Coverage of calcium-related gene ontology enrichment (molecular function). GOMF, KEGG and Uniprot Keyword annotations were used for enrichment analysis using DAVID informatic platforms and Reactome and required a minimum category size of at least three proteins. Percentages of coverage in the indicated MF GO categories are shown. KEGG pathway analysis, were applied to the data, identifying members of major biological processes including muscle contraction and adhesion related (focal adhesion, tight junction) pathways, with percentages of pathway coverage in the indicated KEGG categories shown. Data were subsequently also plugged into Qiagen’s ingenuity pathway analysis (IPA) to predict differences in functional protein signaling networks. A 75% confidence interval was used for prediction analysis for two independent proteomic experiments. Next, IPA software compared the two independent results and determined calcium signaling as the most significantly altered pathway (95% confidence interval, BH corrected *p* value).

### Data analysis

Data was analyzed as described in figure legends. Typically, this involved parametric hypothesis testing via *t* tests and ANOVA. Data were corrected for all relevant comparisons, as outlined in-text, and typically comprised Bonferroni or Tukey’s adjustment. Data were typically represented as mean ± SEM. Relevant variations to analysis (e.g., calcium imaging and proteomics) are described in the relevant section of methods.

## Acknowledgements

We thank the WCM Proteomics Core facility for providing experimental consultation and results. Especially, Dr. Guoan Zhang, the director, and two associate members, Mengmeng Zhu and Taojunfeng Su. We also thank the WCM Genomics Resources Core Facility for consultation and WES sequencing. Additionally, we thank Dr. Zhengming Chen, Ph.D., M.P.H., M.S., a senior research biostatistician in the Division of Biostatistics and Epidemiology at Weill Cornell Medicine, for statistical power analysis. We would like to thank Dr. Lavo Ramos-Espiritu and the High Throughput and Spectroscopy Resource Center at Rockefeller University for training on plate reader for Ca^2^ imaging experiments and valuable comments. This work was supported by a NIH grant 1R01MH120156-01 to D.C.

## Author contributions

D.C. and M.A. conceived of the project, designed most of the experiments and wrote the manuscript with input from all authors. B.S.H. designed and performed live-animal Ca^2+^ imaging experiments, analyzed the data, and helped with the manuscript. P.W. analyzed the WES data and prepared the Supplementary Tables 2 and 3 with input from M.E.R. M.J.N. assisted with 3D cerebral organoid cultures and performed all fear-conditioning and marble burying tests with the help of E.B.A. and A.L. M.A. performed co-culture and other behavioral experiments. M.A., A.L. and E.B.A. executed iPSC and murine cell culture experiments, immunohistochemical experiments, and associated analyses. M.A. and J.W. performed *in vitro* Ca^2+^ imaging experiments with input from C.L. F.L. performed and analyzed LTP experiments with input from E.K. D.G. and M.C. performed proteomics analysis.

## Declaration of interests

The authors declare no competing interests.

## Notes

### Competing Interest Statement

The authors have declared no competing interest.

