## Supplementary figures, legends and tables for "Astrocytes derived from ASD patients alter behavior and destabilize neuronal activity through aberrant Ca^2+^ signaling"

##### Supplementary Information

Supplementary information includes 4 tables, 12 Figures, and 3 videos.

#### Supplementary Tables

**Supplementary Table 1: Information about CTRL subjects and ASD patient lines**

**Supplementary Table 2: Counts of hits for LGD SNVs and LGD CNVs in CTRL subjects and ASD patients**

**Supplementary Table 3: Variants in CTRL and ASD cases and which individual carries them**

**Supplementary Table 4: CTRL subject and ASD patient lines used in each experiment**

#### Supplementary Figures

**Supplementary Fig. 1, Related to Fig. 1:** Organoid-derived astrocytes express astrocyte markers.

**Supplementary Fig. 2, Related to Fig. 1:** Astrocytes dissociated from organoids and maintained in astrocyte selection media are not overtly reactive.

**Supplementary Fig. 3, Related to Fig. 1:** Detection of astrocyte specific proteins through proteomic analysis.

**Supplementary Fig. 4, Related to Fig. 2:** No significant difference in human astrocyte numbers between CTRL and ASD astrocyte chimeric brains.

**Supplementary Fig. 5, Related to Fig. 2:** Transplanted human cells do not terminally differentiate to neurons in chimeric brains.

**Supplementary Fig. 6, Related to Fig. 4:** General motor activity is similar between ASD chimeric and CTRL chimeric mice.

**Supplementary Fig. 7, Related to Fig. 4:** Spatial learning, spatial memory and sociability are not impaired in ASD astrocyte chimeric mice.

**Supplementary Fig. 8, Related to Fig. 5:** Characterization of mouse hippocampal neuronal cultures.

**Supplementary Fig. 9, Related to Discussions:** iPSCs derived from ASD patients and CTRL subjects uniformly convert to NPCs.

**Supplementary Fig. 10, Related to Discussions:** Neural progenitors derived CTRL subject and ASD patient iPSCs terminally differentiate to astrocytes upon transplantation into the mouse brain.

**Supplementary Fig. 11, Related to Discussions:** ASD NPC chimeric mice display impaired associative memory but not learning deficits.

**Supplementary Fig. 12, Related to Discussions:** Glutamate levels are not significantly different between CTRL and ASD astrocytes.

#### Supplementary Videos (Provided as separate files)

**Supplementary Video 1, Related to Fig. 1:** ASD astrocytes respond to stimulation with more intense $Ca^{2+}$ transients compared to CTRL astrocytes.

**Supplementary Video 2, Related to Fig. 5:** ASD astrocyte co-cultures display decreased mean network firing rate when compared to CTRL co-cultures and a control group that contains only dissociated mouse neurons.

**Supplementary Video 3, Related to Fig. 7:** Modulation of ASD astrocyte $Ca^{2+}$ mobilization confers protection against deficits in network firing recorded with MEA.

**Supplementary Table 1: Information about CTRL subjects and ASD patient lines**

| ID | Information |
|---|---|
| CW20008 | ASD; male, Asian, sample collected at age 5, low IQ (68), communication and social deficits (ADOS score 16) as well as stereotyped behaviors and restricted interests (ADOS score 5) |
| CW20142 | ASD; male, Asian, sample collected at age 9, restricted and repetitive behavior (ADOS score 2), communication and social deficits (ADOS score 9) |
| CW20081 | ASD; male, White, sample collected at age 15, low IQ (40), poor eye contact, communication and social deficits (ADOS score 18) as well as stereotyped behaviors and restricted interests (ADOS score 5) |
| CW20083 | ASD; male, White, sample collected at age 22, poor eye contact, severe communication and social deficits (ADOS score 18) repetitive behavior (ADOS score 8) |
| CW60115 | ASD; male, White, sample collected at age 4, poor eye contact, social deficits, stereotyped behaviors and restricted interests (ADOS score 3) |
| MH0148698 | ASD; male, White, low IQ, severely affected (Mariani *et al*., 2015) |
| MH0148713 | ASD; male, White, low IQ, severely affected (Mariani *et al*., 2015) |
| CW20026 | ASD; male, White, sample collected at age 10, low IQ (52), communication and social deficits (ADOS score 19) as well as stereotyped behaviors and restricted interests (ADOS score 3) |
| CW20044 | ASD; male, White, sample collected at age 10, low IQ (40), spinning, severe communication deficits (ADOS score 16) as well as stereotyped behaviors and restricted interests (ADOS score 7) |
| MH0159019 | CTRL, female, Hispanic, sample collected at age 29 |
| MH0159020 | CTRL, male, White, sample collected at age 58 |
| MH0159021 | CTRL, male, Hispanic, sample collected at age 32 |
| MH0174677 | CTRL, male, White, male, white, sample collected at age 9 |
| MH0174679 | CTRL, male, White, male, white, sample collected at age 17 |
| MH0174681 | CTRL, male, White, male, white, sample collected at age 8 |
| MH0174686 | CTRL, male, White, sample collected at age 17 |
| GM23279 | CTRL, female, White, sample collected at age 36 |
| GM23256 | CTRL, male, Asian, sample collected at age 30 |

**Supplementary Table 2: Summary stats for LGD SNVs and LGD CNVs in CTRL subjects and ASD patients**

| | CTRL (9 lines) | ASD (9 lines) |
|---|---|---|
| Rare (< 0.01 MAF), Likely Gene-Disrupting (LGD) SNVs/Indels (322 ASD gene panel) | 37 | 33 |
| Rare (< 0.01 MAF), Likely Gene-Disrupting (LGD) CNVs Overlapping ClinVar P/LP | 31 | 25 |

**Supplementary Table 3: Variants in CTRL (37) and ASD (33) lines and which individual carries them**

| Chromosome and gene | Start | End | Alternate Allele | Sample |
|---|---|---|---|---|
| chr1; RERE | 8656247 | 8656247 | CGGTCC | ASD; CW20026_S55 |
| chr1: TEKT2 | 36088007 | 36088007 | T | ASD; CW20083_S56 |
| chr10; ANK3 | 60042719 | 60042719 | T | ASD; CW20026_S55 |
| chr11; SRPRA | 126265815 | 126265815 | T | ASD; CW60115_S58 |
| chr15; MAP1A | 43525917 | 43525917 | A | ASD; CW20044_S54 |
| chr15; MAP1A | 43527664 | 43527664 | T | ASD; CW20044_S54 |
| chr16; TSC2 | 2053391 | 2053391 | T | ASD; CW20081_S59 |
| chr16; TEKT5 | 10635894 | 10635894 | G | ASD; CW60115_S58 |
| chr16; SETD1A | 30964952 | 30964952 | T | ASD; CW20044_S54 |
| chr17; TEKT1 | 6819266 | 6819266 | C | ASD; CW20026_S55 |
| chr19; ZNF180 | 44476523 | 44476523 | A | ASD; MH0148713_S63 |
| chr2; MYT1L | 1943258 | 1943258 | T | ASD; CW20142_S61 |
| chr2; RNF181 | 85595845 | 85595845 | T | ASD; CW20142_S61 |
| chr2; MBD5 | 148489641 | 148489641 | G | ASD; CW20008_S57 |
| chr3; SETD5 | 9464601 | 9464601 | C | ASD; CW20008_S57 |
| chr3; NBEAL2 | 47004352 | 47004352 | T | ASD; CW20083_S56 |
| chr3; SETD2 | 47121443 | 47121443 | A | ASD; CW20142_S61 |
| chr3; CACNA2D3 | 54891399 | 54891399 | T | ASD; MH0148713_S63 |
| chr4; ANK2 | 113354794 | 113354794 | T | ASD; MH0148713_S63 |
| chr4; NR3C2 | 148154604 | 148154604 | T | ASD; CW20081_S59 |
| chr5; ACTBL2 | 57481701 | 57481701 | T | ASD; CW20142_S61 |
| chr7; RELN | 103539111 | 103539111 | T | ASD; CW20083_S56 |
| chr7; RNF133 | 122698017 | 122698017 | G | ASD; CW20026_S55 |
| chr7; KMT2C | 152205112 | 152205112 | G | ASD; CW60115_S58 |
| chr8; CHD7 | 60865328 | 60865328 | T | ASD; CW20026_S55 |
| chr8; ZNF16 | 144931535 | 144931535 | C | ASD; CW20142_S61 |
| chr1; B4GALT3 | 161173722 | 161173722 | A | ASD; MH0148713_S63 |
| chr16; CDH15 | 89192318 | 89192318 | C | ASD; CW20008_S57 |
| chr17; PIGL | 16317813 | 16317813 | T | ASD; CW20083_S56 |
| chr18; CNDP2 | 74519021 | 74519021 | T | ASD; CW20142_S61 |
| chr19;COLGALT1 | 17577383 | 17577383 | A | ASD; CW20008_S57 |
| chr22; A4GALT | 42693060 | 42693060 | A | ASD; CW20142_S61 |
| chr5; NSUN2 | 6600170 | 6600170 | T | ASD; CW20026_S55 |
| chr20; ARFGEF2 | 49017279 | 49017279 | C | CTRL; MH0159020_S9 |
| chr1; A3GALT2 | 33307191 | 33307191 | T | CTRL; MH0174677_S7 |
| chr1; A3GALT2 | 33307415 | 33307415 | A | CTRL; MH0159019_S5 |
| chr1; DPYD | 97573870 | 97573870 | T | CTRL; MH0174681_S3 |
| chr1; GJA8 | 147908613 | 147908613 | G | CTRL; MH0174681_S3 |
| chr1; PDE4DIP | 149021033 | 149021033 | A | CTRL; GM23279_S2 |
| chr1; ASH1L | 155478027 | 155478027 | C | CTRL; MH0159019_S5 |
| chr1; CACNA1E | 181776219 | 181776219 | A | CTRL; MH0159019_S5 |
| chr1; CACNA1E | 181783712 | 181783712 | A | CTRL; MH0174677_S7 |
| chr10; PCDH15 | 54213995 | 54213995 | A | CTRL; MH0159020_S9 |

| | | | | |
|---|---|---|---|---|
| chr10; ANK3 | 60042719 | 60042719 | T | CTRL; GM23279_S2 |
| chr11; CLP1 | 57659760 | 57659760 | G | CTRL; MH0159019_S5 |
| chr11; SHANK2 | 70490374 | 70490374 | T | CTRL; MH0159019_S5 |
| chr13; NBEA | 35665135 | 35665135 | T | CTRL; MH0174679_S2 |
| chr16; CASKIN1 | 2179616 | 2179616 | A | CTRL; MH0174677_S7 |
| chr16; CASKIN1 | 2181024 | 2181024 | C | CTRL; MH0174677_S7 |
| chr16; TEKT5 | 10694767 | 10694767 | C | CTRL; MH0174681_S3 |
| chr17; RNF135 | 30999137 | 30999137 | T | CTRL; MH0174677_S7 |
| chr18; ASXL3 | 33739587 | 33739587 | A | CTRL; MH0159020_S9 |
| chr19; PRR12 | 49597037 | 49597037 | G | CTRL; MH0159021_S6 |
| chr19; ZNF175 | 51587688 | 51587688 | A | CTRL; GM23256_S8 |
| chr2; BAZ2B | 159412481 | 159412481 | T | CTRL; MH0174679_S2 |
| chr2; SCN2A | 165367225 | 165367225 | T | CTRL; MH0159019_S5 |
| chr2; SCN1A | 166073486 | 166073486 | T | CTRL; MH0174681_S3 |
| chr2; ERFE | 238165677 | 238165677 | G | CTRL; MH0159021_S6 |
| chr3; SETD5 | 9464601 | 9464601 | C | CTRL; GM23256_S8 |
| chr3; SETD2 | 47121346 | 47121346 | C | CTRL; MH0174686_S1 |
| chr3; RNF123 | 49699104 | 49699104 | T | CTRL; GM23256_S8 |
| chr3; PCCB | 136301086 | 136301086 | G | CTRL; MH0159020_S9 |
| chr5; SLC6A18 | 1244318 | 1244318 | T | CTRL; MH0159019_S5 |
| chr5; GABRB2 | 161330900 | 161330900 | A | CTRL; GM23256_S8 |
| chr6; HIVEP2 | 142774335 | 142774335 | C | CTRL; GM23279_S2 |
| chr6; ARID1B | 156778665 | 156778665 | A | CTRL; MH0159019_S5 |
| chrX; NEXMIF | 74740232 | 74740232 | A | CTRL; MH0174686_S1 |
| chrX; PCDH19 | 100296404 | 100296404 | T | CTRL; GM23256_S8 |
| chrX; FLNA | 154353346 | 154353346 | A | CTRL; MH0174686_S1 |
| chr1; RERE | 8361046 | 8361046 | GGGATGCGGCGG | CTRL; MH0159021_S6 |

**Supplementary Table 4: CTRL subject and ASD patient lines used in each experiment**

| Experiment | Lines |
|---|---|
| Whole Exome Sequencing (Supp Table 2 and 3) | All lines in Supp table 1 (9 ASD, 9 CTRL) |
| Proteomics (Fig.1 and Supp Fig. 3) | All lines in Supp table 1 except a CTRL line MH0159019 (9 ASD, 8 CTRL) |
| Astrocyte reactivity (Supp Fig. 2) | All lines in Supp table 1 (9 ASD, 9 CTRL) |
| *In vitro* $Ca^{2+}$ imaging (Fig. 1 and 7) | CTRL: MH0159020, MH0159021, MH0174677 ASD: CW20008, CW20083, CW60115, CW20142, |
| MEA (Fig. 5) | CTRL: MH0159020, MH0159021, MH0159019 ASD: CW20008, CW20083, CW60115, CW20081 |
| Spine quantifications (Fig. 5) | CTRL: MH0159020, MH0159021, MH0174677, MH0159019; ASD: CW20008, CW20083, CW60115, CW20142, CW20081 |
| Stereological Quantifications (Supp Fig. 4) | CTRL: MH0159020, MH0159021, MH0174677 ASD: CW20008, CW20083, CW60115 |
| NeuN+GFP+ co-localization in chimeric brains (Supp Fig. 5) | CTRL: MH0159020, MH0159021, MH0174677, MH0159019; ASD: CW20008, CW20083, CW60115, CW20142, CW20081 |
| Live animal $Ca^{2+}$ imaging (Fig. 3) | CTRL: GM23256, MH0159020, MH0159021, MH0174677; ASD: CW20008, CW20083, CW20142, CW20081 |
| Behavior or LTP experiments in astrocyte chimeric mice (Fig. 4) | CTRL: MH0159019, MH0159020, MH0159021, MH0174677, GM25256; ASD: CW20008, CW20142, CW20081, CW20083, CW60115 |
| MWM or Sociability (Supp Fig. 8) | CTRL: MH0159019, MH0159020, MH0159021, MH0174677; ASD: CW20008, CW20081, CW60115 |
| Rescue- MEA (Fig. 7) | CTRL: MH0159020, MH0159021, MH0174677 ASD: CW20083, CW60115, CW20142, CW20081 |
| Rescue- Fear Conditioning (Fig. 7) | CTRL: MH0159021, MH0159019 ASD: CW20083, CW60115, CW20142, CW20081 |
| NPC experiments (Supp Fig. 9-11) | CTRL: MH0159019, MH0159021, MH0159021; ASD: CW20008, CW60115, CW20142, CW20083 |
| Glutamate Assay (Supp Fig. 12) | All lines in Supp table 1 (9 ASD, 9 CTRL) |

Supplementary Fig. 1

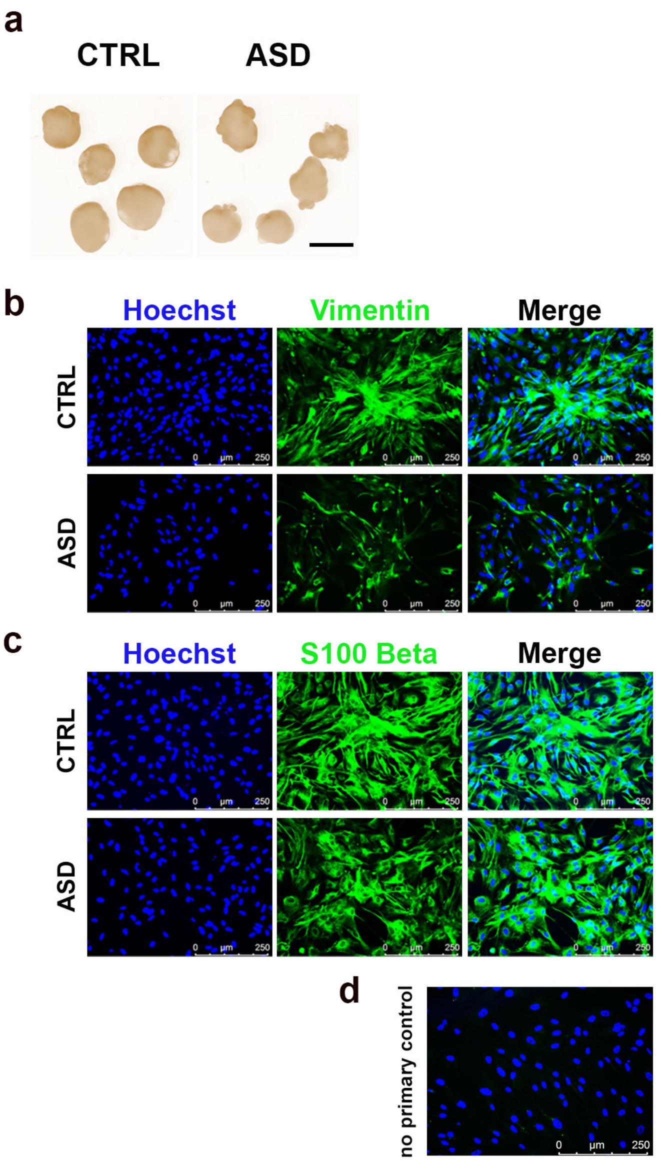


**Supplementary Fig. 1: Organoid-derived astrocytes express astrocyte markers, related to Fig. 1.**

**a,** Representative images of CTRL and ASD cerebral organoids at day 70.

**b,c,** Representative images from immunostainings showed that cells dissociated from organoids (at day 75) expressed Vimentin (**b**) and S100Beta (**c**), two well-described astrocyte markers (see also **Fig. 1** and **Supp Fig. 3** for additional markers) **d,** Representative no primary control immunostained image demonstrated the specificity of positive signal in b and c. No-primary control was used to determine imaging exposure times. Scale bar = 250 μm.

**Supplementary Fig. 2**

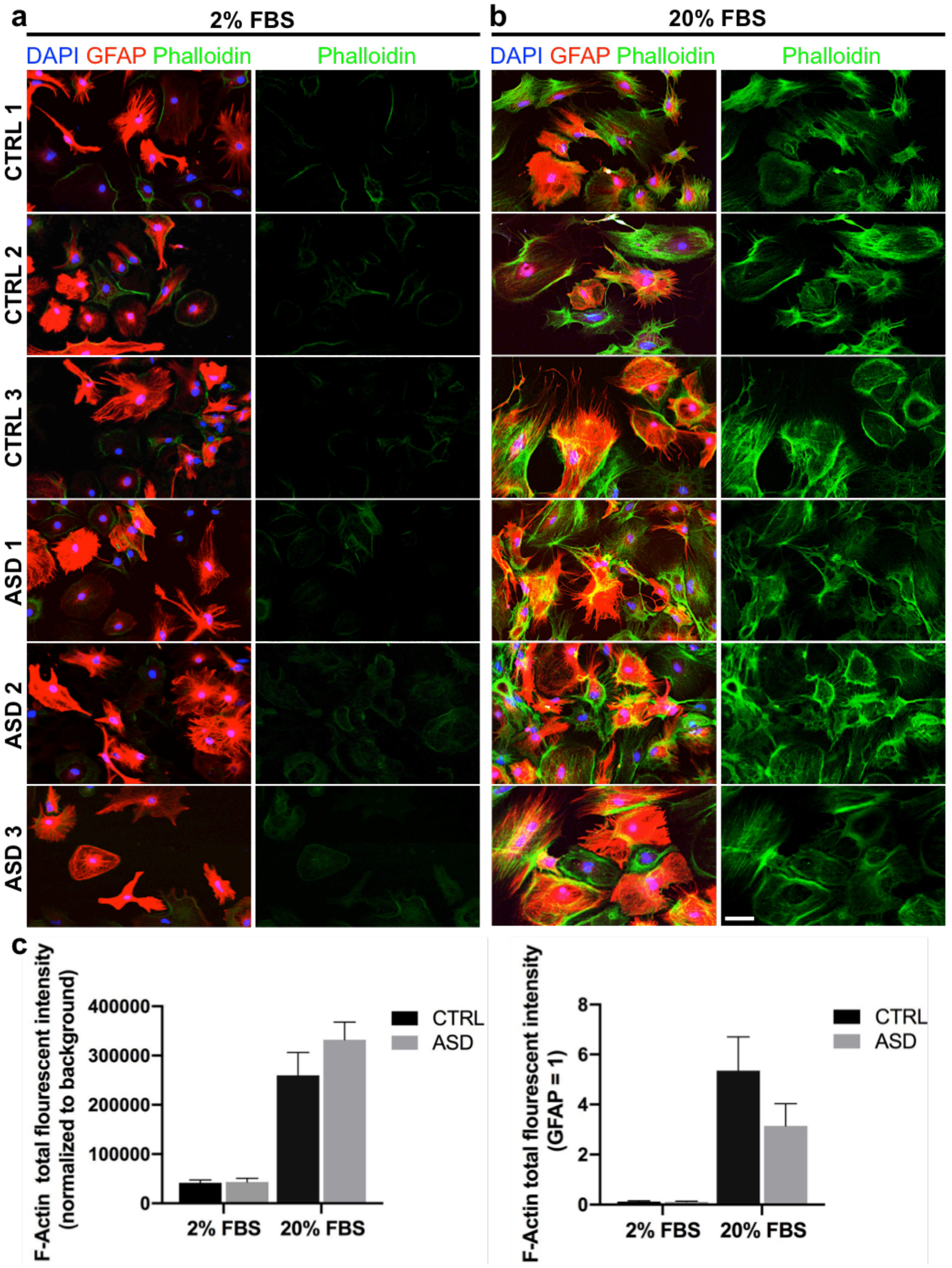

**Supplementary Fig. 2: Astrocytes dissociated from organoids and maintained in astrocyte selection media are not overtly in reactive state, related to Fig. 1.**
To assess whether organoid-derived CTRL or ASD astrocytes were reactive, we visualized F-Actin by Phalloidin labeling and quantified F-Actin fluorescent intensity. Our astrocyte selection media contains minimal FBS (2%) that is required for astrocyte proliferation and known to cause no reactivity in these cells. Because high levels of FBS is known to induce reactivity in astrocytes, we cultured both CTRL and ASD organoid derived astrocytes in astrocyte selection media with 20% FBS, as a positive control, only in this experiment. Quantifications in **c** were done in astrocytes generated from all CTRL (9 distinct iPSC lines) and ASD (9 distinct iPSC lines) lines used in this paper.

In addition, we also assessed the reactivity status of our astrocytes by screening for specific reactive astrocyte markers within our astrocyte proteomics datasets. For this, we consulted two papers[1, 2] that together identified 12 markers whose levels were increased only when astrocytes are reactive. Of these, 9 reactive astrocyte markers *were not detected* via TMT-LC/MS proteomics in any of our CTRL or ASD samples (SERPINA3N, TNFRSF12A, S1PR3, CXCL1, CXCL2, CXCL10, LCN2, STEAP4, and OSMR). 3 reactive astrocyte markers (PTX3, TIMP1, and CD44) maintain other homeostatic cellular activities and were identified in our samples as expected. However, expression levels of these remaining factors were relatively low in abundance and without any significant p value or Log2FC value change.

**a,b,** Representative images of astrocytes (isolated from organoids derived from *n* = 3 distinct iPSCs per group) that were cultured in 2% (**a**) or 20% (**b**) FBS, and stained with GFAP and Phalloidin. While F-Actin signal can be seen at similar and relatively low levels in both CTRL and ASD astrocytes in our regular culture conditions (2% FBS), the F-Actin signal is highly increased in these cells when cultured in reactivity-inducing conditions, i.e., 20% FBS. **c,** We measured total F-Actin fluorescent in astrocytes generated from all CTRL and ASD iPSC lines used in the paper (*n* = 9 distinct ASD lines, *n* = 9 distinct CTRL lines). In both quantifications when either normalized to background (left) or, as an internal control, to GFAP (right), F-Actin levels in both group found to be significantly increased in reactivity-inducing conditions.

**Together, these data suggest that CTRL and ASD organoid-derived astrocytes that are maintained in astrocyte selection media are not in a reactive state.**
Scale bar = 50 μm Data are represented as mean ± SEM. *n* = 9 distinct ASD lines, *n* = 9 distinct CTRL lines.

### Supplementary Fig. 3

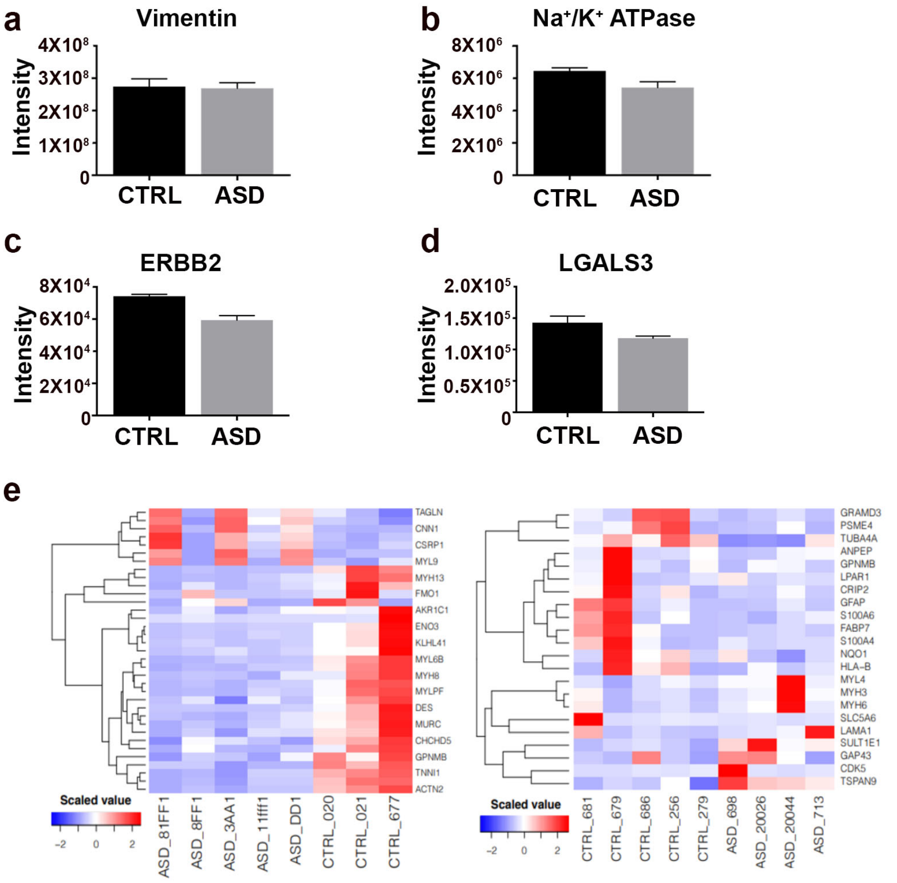


**f**

| Protein IDs | Gene names | Protein names | log2 FC (ASD/Ctrl) | p-value | Regulation (ASD/Ctrl) |
|---|---|---|---|---|---|
| Q96A32 | MYLPF | Myosin regulatory light chain 2, skeletal muscle isoform | -3.555 | 1.25E-07 | down-regulated |
| P63316 | TNNC1 | Troponin C, slow skeletal and cardiac muscles | -2.953 | 7.67E-06 | down-regulated |
| P35609 | ACTN2 | Alpha-actinin-2 | -2.001 | 6.26E-03 | down-regulated |
| P17302 | GJA1 | Gap junction alpha-1 protein | -1.918 | 8.81E-03 | down-regulated |
| P19237 | TNNI1 | Troponin I, slow skeletal muscle | -1.893 | 1.29E-02 | down-regulated |
| P21980 | TGM2 | Protein-glutamine gamma-glutamyltransferase 2 | 1.534 | 3.24E-02 | up-regulated |
| Q01995 | TAGLN | Transgelin | 1.788 | 6.15E-03 | up-regulated |
| P24844 | MYL9 | Myosin regulatory light polypeptide 9 | 1.965 | 1.60E-03 | up-regulated |

| Protein IDs | Gene name | Protein names | log2 FC (ASD/Ctrl) | p-value | Regulation (ASD/Ctrl) |
|---|---|---|---|---|---|
| P06703 | S100A6 | Protein S100;Protein S100-A6 | -2.407 | 3.15E-05 | down-regulated |
| P26447 | S100A4 | Protein S100-A4 | -1.965 | 9.00E-04 | down-regulated |
| Q92633 | LPAR1 | Lysophosphatidic acid receptor 1 | -1.561 | 2.22E-02 | down-regulated |
| Q00535 | CDK5 | Cyclin-dependent-like kinase 5 | 2.005 | 1.24E-03 | up-regulated |

**Supplementary Fig. 3: Detection of astrocyte specific proteins through proteomic analysis, related to Fig. 1.**

To identify the inherent pathological properties within ASD astrocytes, we took an unbiased approach and compared the protein profiles of 9 ASD patient astrocytes to 8 healthy CTRL subject astrocytes with Tandem mass tag (TMT) liquid chromatography mass spectrometry.

**a-d,** Proteomic analysis detected several markers for astrocyte identity in our samples and the intensities are compared in the graphs.

**e,f,** Expression based heat maps from two independent experiments exhibited quantitative patterns across proteins and biological samples (CTRL $n$ = 8 unique patients and ASD $n$ = 9 unique patients). Each column represents astrocytes from a unique patient, while relevant proteins are represented in each row. (BH adjusted $p$-values > 0.05). Due to statistical correctness, these independent data cannot be pooled together as the experiments and analyses were not performed at the same time (as part of the same TMT-plex, due to the limitations in how many samples could be barcoded and run simultaneously at the time experiments were performed).

**Supplementary Fig. 4**

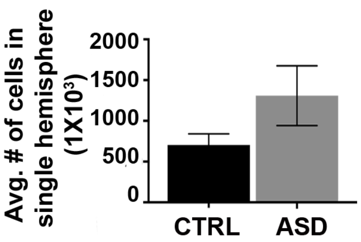


**Supplementary Fig. 4: No significant difference in human astrocyte numbers between CTRL and ASD astrocyte chimeric brains, related to Fig. 2.**
We dissociated astrocytes from CTRL or ASD cerebral organoids at day 75 and expanded in 2D adherent culture. Astrocytes were labeled with a lentivirus expressing GFP prior to transplantation into *Rag2*$^{KO}$ postnatal mouse brains between P1-3. To assess the average number of astrocytes that survived in host mice, we performed unbiased stereological cell counting on GFP immunostained brain sections. We serially sectioned chimeric brains at P60 along the sagittal plane (30 μm thickness). Every fourth section was used for analysis. Using the optional fractionator method, we estimated the number of surviving cells in one hemisphere from mice transplanted 3 distinct lines for CTRL or ASD. We did not find a significant difference between the numbers of human astrocytes in hemisected CTRL or ASD astrocyte chimeric brains (CTRL: 704,684 ±135,971 GFP+ cells, ASD: 1,309,082 ±367,310 GFP+ cells, unpaired $t$ test $p$ value = 0.21). CTRL $n$ = 3 transplanted brains, 3 distinct lines, ASD $n$ = 3 transplanted brains, 3 distinct lines.

Supplementary Fig. 5

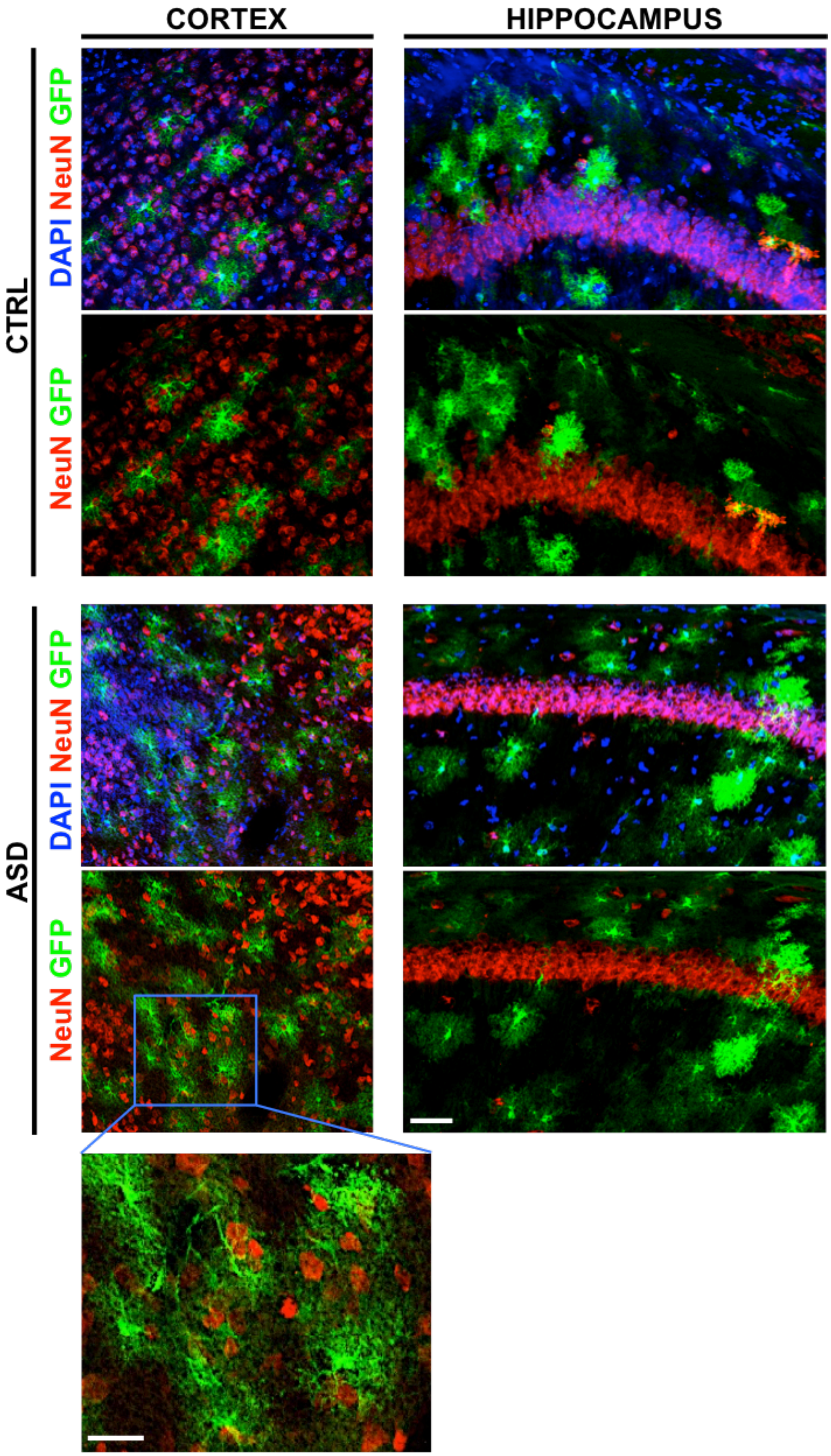

**Supplementary Fig. 5: Transplanted human cells do not terminally differentiate to neurons in chimeric brains, related to Fig. 2.**

We transplanted healthy CTRL or ASD astrocytes into the brains of *Rag2*$^{KO}$ mice between postnatal days 1-3 (P1-3). **Fig. 2** shows that more than 90% of the transplanted cells co-localize with known astrocyte markers in adult chimeric brains. Here, we present representative images showing that none of the transplanted GFP+ cells co-localized with the neuronal marker NeuN. We examined over 800 GFP+ cells in 9 chimeric brains (transplanted with 4 distinct iPSC lines/4 CTRL subjects and 5 distinct iPSC lines/5 ASD patients; same lines used in behavioral experiments in **Fig. 4**) and found that none of these GFP+ cells expressed nuclear NeuN or exhibited a typical neuronal morphology. Scale bar = 100 μm for all low magnification panels and 50 μm for the high magnification image at the bottom.

**Supplementary Fig. 6**

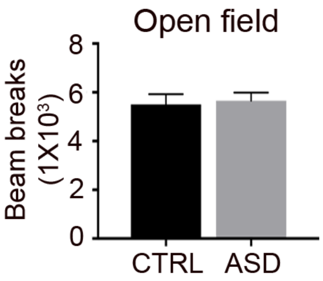


**Supplementary Fig. 6: General motor activity is similar between CTRL chimeric and ASD chimeric mice, related to Fig. 4.**

We evaluated general locomotor activity in chimeric mice with an automated open field test. ASD chimeric mice did not demonstrate differences in exploratory or general activity relative to CTRL chimeric mice (CTRL 5502 ±423.6 beam breaks, ASD 5659 ±334 beam breaks, unpaired $t$ test, $p$ value = 0.782). These results suggest that our injection protocol did not affect ambulatory activity and is not a confounding factor in the interpretation of other behavioral results. CTRL $n$ = 22 mice (3 distinct lines), ASD $n$ = 17 mice (3 distinct lines). Data are represented as mean ± SEM.

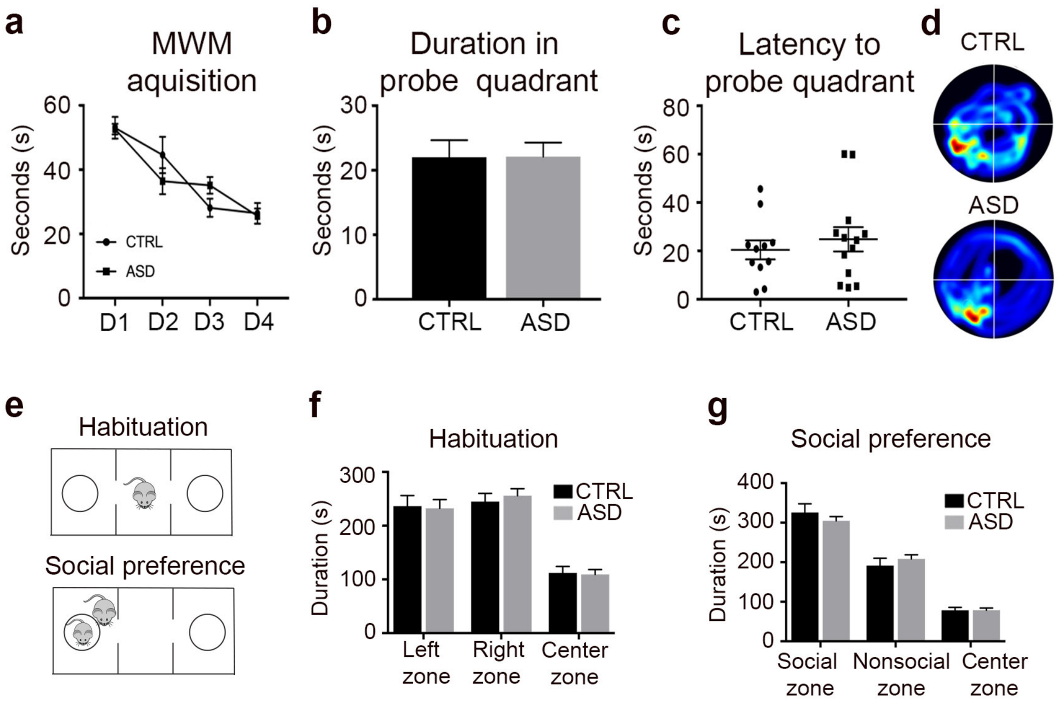


**Supplementary Fig. 7: Spatial learning, spatial memory and sociability are not impaired in ASD astrocyte chimeric mice, related to Fig. 4.**

**a-d,** ASD astrocyte chimeric mice showed no deficits in spatial learning or memory when compared to CTRL. To test spatial memory, we used the Morris Water Maze (WMW), where a test mouse relies on distal cues to locate a hidden escape platform submerged in opaque water. Spatial learning is acquired though repeated trials and memory is determined through efficient navigation to an area that formerly housed the escape platform. **a**, We found no difference between CTRL and ASD mice in the latency to locate the hidden platform during training, which consisted of 4 trials over 4 consecutive days (ANOVA $p$ value > 0.05). **b**,**c,** On the fifth day, the hidden platform was removed and the duration (s) spent in the quadrant that formerly housed the platform was measured. No differences between the latency (s) to reach the correct quadrant were detected between CTRL and ASD (CTRL: 20 ±3.91s and ASD 25 ±5.02s, unpaired $t$ test = 0.51). **d**, Representative heat maps of mouse movements CTRL $n$ = 11 male and female mice, transplanted with 3 distinct lines, ASD $n$ = 13 male and female mice, transplanted with 3 distinct lines.

**e-g,** To assess sociability, we administered 3-chambered sociability assay in astrocyte chimeric mice (**e**). **f,** During the habituation phase, both CTRL and ASD astrocyte chimeric mice explored the left, right, and center chambers for similar durations suggesting that no bias existed before the insertion of a social stimulus (Left: CTRL

236.35 ±19.76s and ASD 232.41 ±16.13s, Right: 244.82 ±15.24s and ASD 255.77 ±13.18s, Center: CTRL 112.08 ±12.10s and ASD 109.30 ±9.11s, ANOVA with Bonferroni posttests, $p$ value > 0.05). **g**, ASD astrocyte chimeric mice spent a relatively similar amount of time in the social chamber that contained the stimulus mouse compared to CTRL (Social chamber: CTRL 325.74 ±22.14s and ASD 304.71 ±10.77s, Nonsocial chamber: CTRL 191.61 ±18.44s and ASD 208.41 ±10.35s, Center CTRL 78.60 ±7.28s and ASD 78.76 ±5.52s, ANOVA with Bonferroni posttests $p$ value > 0.05). CTRL $n$ = 12 male and female mice, transplanted with 3 distinct lines, ASD $n$ = 22 male and female mice, transplanted with 3 distinct lines. Data are represented as mean ± SEM.

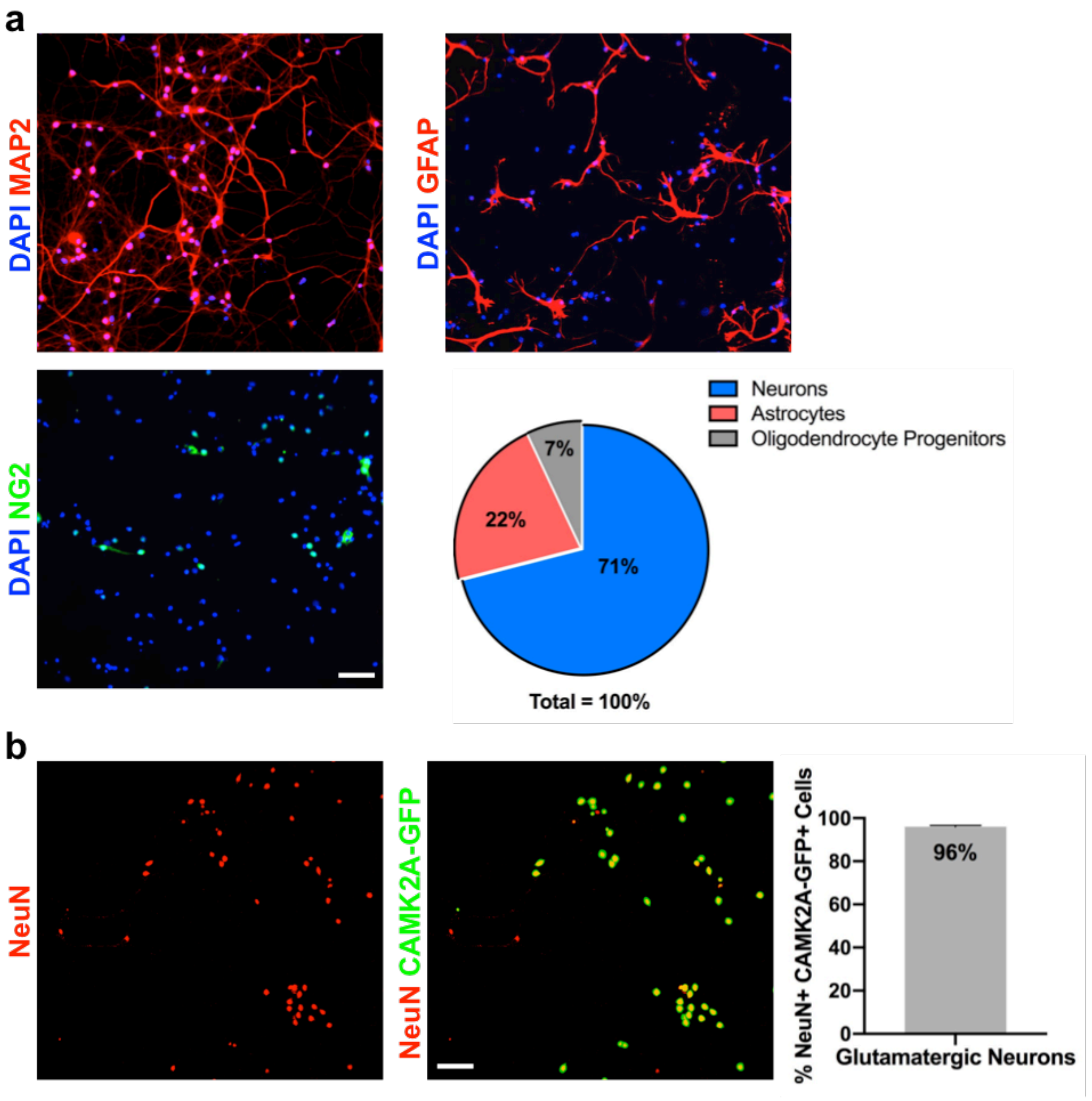


**Supplementary Fig. 8: Characterization of mouse hippocampal neuronal cultures, related to Fig. 5.** We dissociated primary hippocampal neurons from the hippocampi of C57BL/6N embryos at E16-18 as previously described[3] and plated $1x10^5$ cells. **a,** At DIV 18, more than 70% of the cultured cells acquired neuron fate, while astrocytes and other glial progenitors represented 22% and 7% of cells, respectively. **b,** Infection of these cultures with a GFP reporter in which nuclear GFP is driven from CAMK2A promoter revealed that more than 95% of the neurons are excitatory glutamatergic neurons.

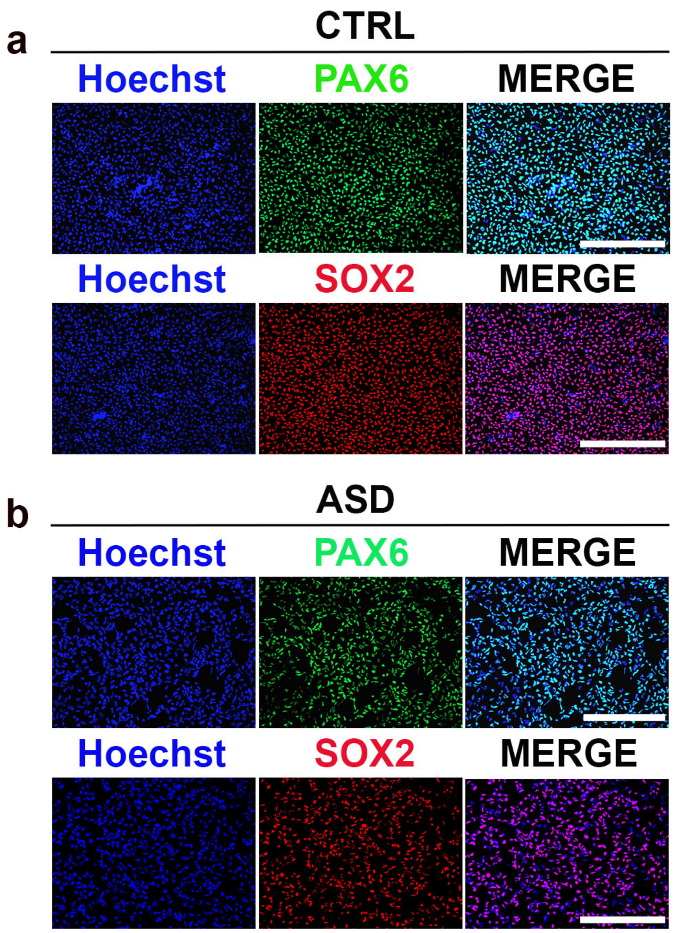


**Supplementary Fig. 9: iPSCs derived from ASD patients and CTRL subjects uniformly convert to NPCs, related to Discussions.**

**a,b,** To induce differentiation of iPSCs to forebrain Neural Progenitor Cells (NPCs), we used an established dual SMAD inhibition protocol[4]. Representative images of iPSC-derived NPCs from CTRL subjects (**a**) and ASD patients (**b**) highlighted homogenous conversion with the dual SMAD inhibition protocol as evidenced by expression of PAX6 and SOX2, two early neural progenitor markers. Scale bar = 250 μm.

Supplementary Fig. 10

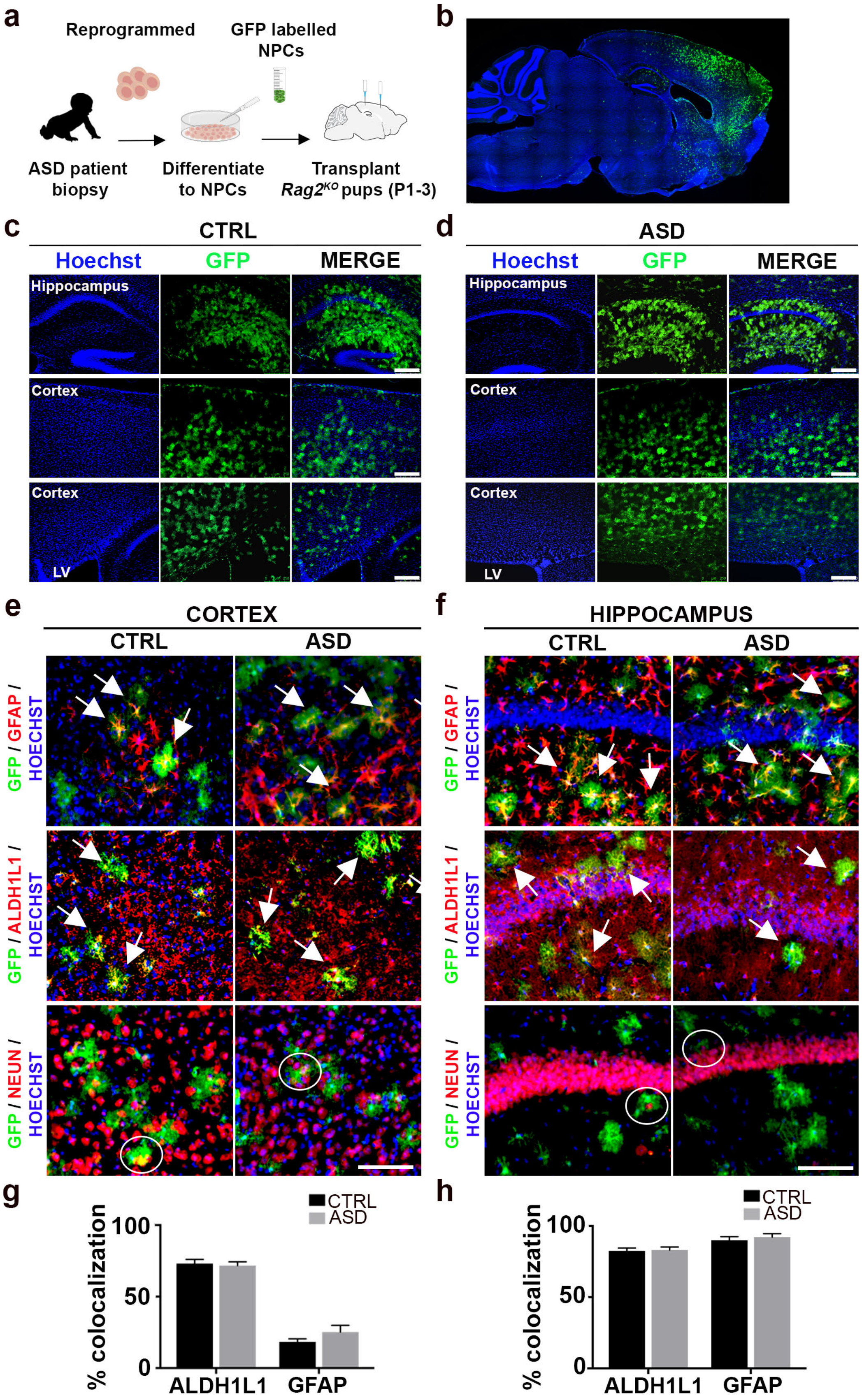

**Supplementary Fig. 10: Neural progenitors derived from CTRL and ASD iPSC lines terminally differentiate to astrocytes upon transplantation into the mouse brain, related to Discussions.**

We transplanted healthy CTRL or ASD neural progenitor cells (NPCs) into the brains of immunocompromised $Rag2^{KO}$ mice between postnatal days 1-3 (P1-3). In the healthy developing mouse brain, extensive gliogenesis typically occurs between embryonic day 17 and P17, with peak astrocyte formation occurring between P0 and P2 and oligodendrocyte formation at P14[5]. To induce differentiation of iPSCs to early NPCs, we used an established dual SMAD inhibition protocol[4] (see also **Supp Fig. 9**). Prior to transplantation, human NPCs were labeled with a CAG-GFP lentivirus for detection.

**a,** Schematic summarizing the experimental workflow. All immunostainings were performed at P60. **b-d,** We transplanted GFP+ human NPCs into $Rag2^{KO}$ mouse brains at P1-3. To visualize the transplantations in the brain, we immunostained whole brain slices cut on the sagittal plane with GFP (**b**, a representative ASD NPC chimeric section). We found no gross differences in the migration or distribution of human cells in ASD or CTRL NPC chimeric brains. **c**,**d,** Representative higher magnification images displayed the extent of human GFP+ cell infiltration throughout the somatosensory cortex and hippocampus in CTRL and ASD transplanted brains. **e**-**h,** GFP+ cells displayed an astrocyte-like morphology in the mature mouse brain. To quantify the number of NPCs that terminally differentiated to astrocytes, we coimmunostained chimeric mouse brains for GFP and either GFAP or ALDH1L1, two well described astrocyte markers. To rule out differentiation to neuronal cell fate, we also coimmunostained chimeric brains for GFP and the neuronal marker, NeuN. Representative coimmunostained images of chimeric brains revealed colocalization of GFP expression (human cells, green) with the expression of two astrocyte markers, GFAP (red, top panel) and ALDH1L1 (red, middle panel) in the cortex (**e**) and hippocampus (**f**). Arrows indicate examples of dual positive cells. Notably, we did not find colocalized expression of GFP (human cells, green) and NeuN (red, bottom panel). However, we observed GFP+ processes cradling mouse NeuN positive cell bodies (empty circles). **g**,**h,** More than 70% of GFP expression colocalized with the expression of astrocyte markers both in cortex (**g**) and hippocampus (**h**) (Cortex: ALDH1L1/GFP CTRL 73% ±3% and ASD 72% ±3%, GFAP/GFP CTRL 18% ±2% and ASD 25% ±5%; Hippocampus: ALDH1L1/GFP HP: CTRL 82% ±2% and ASD 83% ±2%, GFAP/GFP CTRL 90% ±3% and ASD 92% ±2%).

**Results from these experiments confirm that CTRL and ASD NPCs generated homogenous, high-density transplantations in host brains and primarily differentiated into astrocytes upon maturation in the living brain.**

Scale bar = 250 μm for c-f. Data are represented as mean ± SEM. Immunostaining quantifications: CTRL and ASD *n* = 12 per group (3 brains transplanted with distinct lines per group and 4 slices per brain). Data are represented as mean ± SEM.

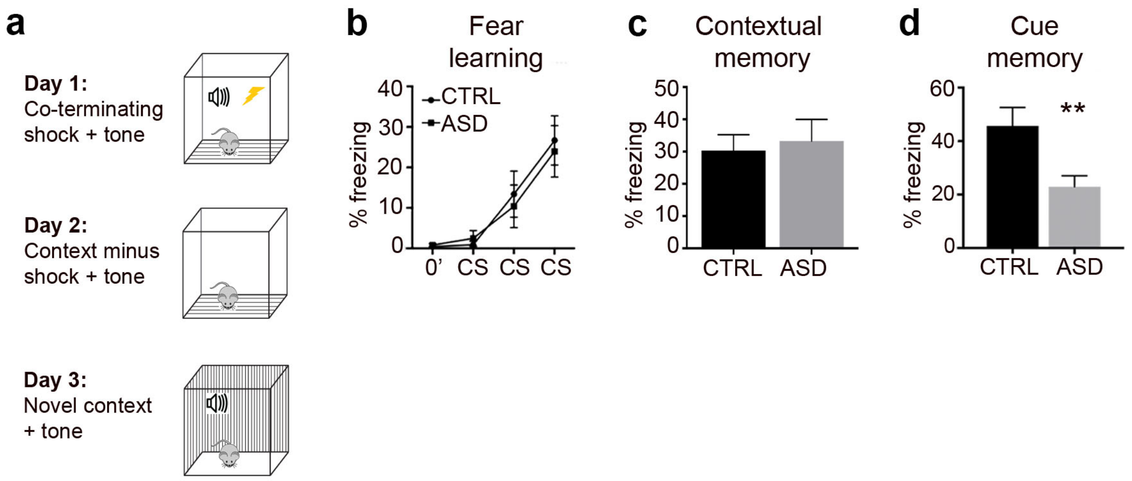


**Supplementary Fig. 11: ASD NPC chimeric mice display impaired associative memory but not learning deficits, related to Discussions.**
We transplanted healthy control or patient neural progenitor cells (NPCs) into the brains of immunocompromised *Rag2*$^{KO}$ mice between postnatal days 1-3 (P1-3). In the healthy developing mouse brain, extensive gliogenesis typically occurs between embryonic day 17 and P17, with peak astrocyte formation occurring between P0 and P2 and oligodendrocyte formation at P14[5] (see also **Supplementary Fig. 9** for NPC production). Greater than 70 percent of transplanted human CTRL or ASD NPCs acquired an astrocytic fate *in vivo* (**Supplementary Fig. 10**).

**a-d,** We tested NPC chimeric mice for deficits in learning and memory with a classical fear conditioning protocol. Fear conditioning to a cued signal or a specific context is a form of associative learning and memory. In this paradigm, freezing behavior represented a species-specific response to fear and was defined as the absence of movement except for respiration. **a,** Schematic summarizing classical fear conditioning paradigm. On day 1, mice learned to associate an audible tone (30 sec duration, 70dB) with a co-terminating foot shock (1 sec duration, 0.7mA). Testing days 2 and 3 measured freezing behavior in response to exposure to the training context or the audible cue in a novel context, respectively. Freezing behavior during these testing trials provided a quantifiable measure of associative memory. **b,** Both CTRL and ASD NPC chimeric mice learned the association as evidenced by increased freezing behavior in response to each successive tone-shock pairing (ANOVA with Bonferroni posttests *p* value = 0.0001). We found no significant differences in the rate of learning between CTRL and ASD chimeric mice (ANOVA with Bonferroni posttests *p* value > 0.05).

**c,d,** Transplantation of ASD NPCs did not influence contextual associative learning (CTRL 30.33 ±4.91%, ASD 33.38 ±6.69%, unpaired *t* test, *p* value = 0.73) (**c**). However, ASD NPC chimeric mice displayed lower freezing levels than CTRL during the cued memory trial indicating impaired associative memory (CTRL 45.68 ±6.89%, ASD 22.95 ±4.07%, unpaired *t* test, *p* value = 0.007) (**d**). CTRL n = 12 mice, transplanted with 3 distinct lines, ASD n = 14 mice, transplanted with 4 distinct lines. Data are represented as mean ± SEM.

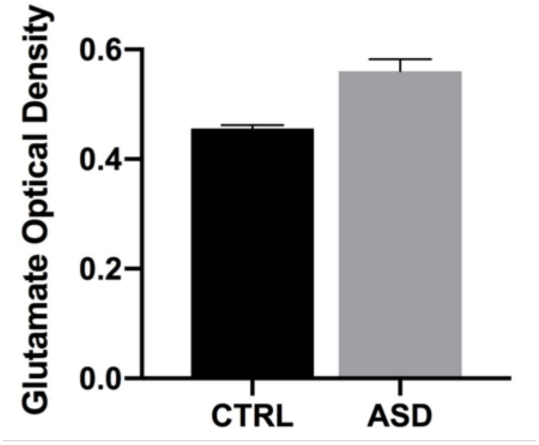


**Supplementary Fig. 12: Glutamate levels are not significantly different between CTRL and ASD astrocytes, related to Discussions.** We measured glutamate levels in CTRL and ASD organoid-derived astrocyte supernatant (Passage 8) using a glutamate assay kit (ab83389) (see Methods section). Supernatant was collected 96 hours after passaging and showed no difference in glutamate levels between CTRL and ASD astrocytes ($p$ value: 0.16; $n$ = 9 distinct lines for ASD and $n$ = 9 distinct lines for CTRL). Data are represented as mean ± SEM.

**Supplementary Video 1**, Related to Fig. 1: ASD astrocytes respond to stimulation with more intense $Ca^{2+}$ transients compared to CTRL astrocytes.
To acquire $Ca^{2+}$ signals, we first loaded astrocytes with a $Ca^{2+}$ indicator dye (Fluo-4-am, 1μM) and incubated for 30 min at 37°C. The dye was completely washed away and cells permitted to equilibrate at 37°C for 30 min. We used Olympus RS equipped with a scanning galvanometer and a Spectra-Physics Mai Tai DeepSee laser for the excitation of GFP. We recorded fluorescence through gallium arsenide phosphide (GaAsP) detectors using the Fluoview acquisition software (Olympus) with a green light emission bandpass filter (Semrock).

**Supplementary Video 2**, Related to Fig. 5: MEA recordings from DIV14 human astrocyte and mouse neuron co-cultures and mouse neurons only control culture. Each column represented an astrocyte derived from a unique patient (CTRL *n*=3 and ASD *n*=4). Heat map of real-time firing visually showed that CTRL co-cultures had more spontaneous network activity than ASD co-cultures.

**Supplementary Video 3**, Related to Fig. 7: MEA recordings from DIV14 human astrocyte and mouse neuron co-cultures. Human astrocytes were transduced control (non-targeting, Non) or ITPR knockdown (KD) shRNA and then added to culture with mouse primary neurons. Shown are CTRL Non co-cultures with astrocytes derived from two unique patients. ASD co-cultures were grouped by unique patient. KD treatment in ASD co-cultures increased neuronal network activity so that there was no significant difference when compared to the activity of CTRL Non co-cultures.
